# Time-resolved cryo-EM (TRCEM) sample preparation using a PDMS-based microfluidic chip assembly

**DOI:** 10.1101/2024.12.08.627437

**Authors:** Xiangsong Feng, Joachim Frank

## Abstract

Time-resolved cryo-EM (TRCEM) makes it possible to provide structural and kinetic information on a reaction of biomolecules before the equilibrium is reached. Several TRCEM methods have been developed in the past to obtain key insights into the mechanism of action of molecules and molecular machines on the time scale of tens to hundreds of milliseconds, which is unattainable by the normal blotting method. Here we present our TRCEM setup utilizing a polydimethylsiloxane (PDMS)-based microfluidics chip assembly, comprising three components: a PDMS-based, internally SiO_2_-coated micromixer, a glass-capillary microreactor, and a PDMS-based microsprayer for depositing the reaction product onto the EM grid. As we have demonstrated in recent experiments, this setup is capable of addressing problems of severe sample adsorption and ineffective mixing of fluids, and leads to highly reproducible results in applications to the study of translation. As an example, we used our TRCEM sample preparation method to investigate the molecular mechanism of ribosome recycling mediated by <u>H</u>igh frequency of lysogenization <u>X</u> (HflX), which demonstrated the efficacy of the TRCEM device and its capability to yield biologically significant, reproducible information. This protocol has promise to provide structural and kinetic information on pre-equilibrium intermediates in the 10-1000 ms time range in applications to many other biological systems.

**Key features:**

- Design and fabrication of high-performance splitting-and-recombination-based micromixer and planar microsprayer.
- Protocol for SiO_2_ coating on the PDMS surface and fabrication of the microfluidic chip assembly.
- Preparation of time-resolved cryo-EM sample in the time range of 10 – 1000 ms.
- Data collection on EM grid covered with droplets from the microsprayer.

## Background

Time-resolved cryo-electron microscopy (TRCEM)[1] is on the way to become a key technique to unlock the dynamics of biomolecular reactions by providing occupancies (i.e., kinetic data) and structural manifestations of intermediate states in the approximate time range of 10 to 1000 ms. The setup of a TRCEM experiment must ensure sufficiently thorough mixing of the reactants, precise control of reaction time, and fast vitrification, such that reaction intermediates can be trapped and subsequently visualized by single-particle cryo-EM.

Over the past two decades, many TRCEM methods have been developed[2–10]. These can be grouped into two main categories, spraying/mixing and mixing/spraying. In the first category, a sprayer or dispenser device deposits a reactant onto an EM grid that is pre-covered with another reactant. Both mixing and reacting occur on the EM grid, which is then rapidly plunged into the cryogen for fast vitrification. In this case, the length of the reaction time is controlled solely by the duration of the plunging. However, the uniformity of the reaction is not ensured as it relies on the efficiency of diffusive mixing on the grid and is affected by uncontrolled processes at the air-water interface. In the second category, a microfluidic chip combining the functions of mixing, reacting and spraying is used for controlling the reaction and the deposition of the reaction product onto the grid. Here the plunging time is kept very short to keep the on-grid time to a minimum. Most importantly, mixing and reacting can be separately manipulated, and the reaction time is defined mainly by the residence time of the sample in the microfluidic chip.

For the reasons stated, we have opted for the mixing/spraying, microfluidics-based sample preparation method in our development of the TRCEM technology. This development was driven by the need for migrating from silicon, in the initial version of the microfluidic chip[8], to plastic polymers as more versatile and cost-effective material for the fabrication of microfluidic chips[4,5,10]. However, this migration faced an important obstacle since polymers such as IP-S and IP-Q photoresins and PDMS are intrinsically hydrophobic and prone to adsorption of biomolecular samples, thereby impairing the control over the reactions. Another problem we needed to address – this one shared with silicon-based chips – was ineffective mixing of fluids due to limited micromixer performance in the laminar flow regime.

In our attempt to overcome these problems we upgraded our TRCEM setup with a PDMS-based microfluidic chip assembly comprising three modules [11]: 1) a PDMS-based splitting-and-recombination-based (SAR) micromixer with 3D self-crossing channels, which is able to efficiently mix the solutions for uniform initiation of a reaction; 2) a polyimide-coated glass capillary tubing, which ensures precise flow-rate control of liquids, to serve as the microreactor whose dimensions define the reaction time; 3) a PDMS-based microsprayer (with a design improved from our previous microsprayer [12]) employed to spray out the droplets of the reaction product onto the EM grid. This sprayer has been demonstrated in our work to be efficient for preparing cryo-EM grids with vitreous ice of controllable, highly consistent thickness. The details of fabrication of these three modules, their integration into a functioning assembly mounted in the TRCEM apparatus, and the testing of the entire setup with biological samples are described in the following.

We believe that the protocol contains sufficient detail to allow other teams with appropriate scientific and engineering skills to fabricate the microfluidic chips, assemble the entire functioning apparatus, and duplicate the experiments described.

## Materials and reagents

### Biological materials

Note: Since this protocol is focused on the preparation of TRCEM grids, the purification of biological materials (*E. coli* 70S ribosome and *HflX*) will not be explained here. Note that HflX is known as a universally conserved protein for prokaryotic cells, a GTPase that functions as a rescue factor which splits the 70S ribosome into its 30 S and 50S subunits in response to heat shock or exposure to antibiotics. For details, the reader is referred to ref.[11] and literature cited therein.

1. *E. coli* 70S ribosome (1 mM in the buffer solution)
2. *HflX* (5 mM in the buffer solution)

### Reagents

1. GTP (Invitrogen Cat# 18332015).
2. Methylene blue (Sigma-Aldrich Cat# 457250)
3. Tris Base (Fisher Scientific Cat# BP152-1)
4. Ammonium chloride (NH_4_Cl) (Sigma-Aldrich Cat# 09718)
5. Magnesium acetate (Mg(OAc)_2_) (Sigma-Aldrich Cat# M5661)
6. 2-Mercaptoethanol (βME) (Sigma-Aldrich Cat# M6250)

### Solutions

1. Methylene blue solution (see Recipes).
2. Buffer solution (see Recipes).

### Recipes

1. **Methylene blue solution (10 mL) [Used within 3 years, and stored at room temperature]** 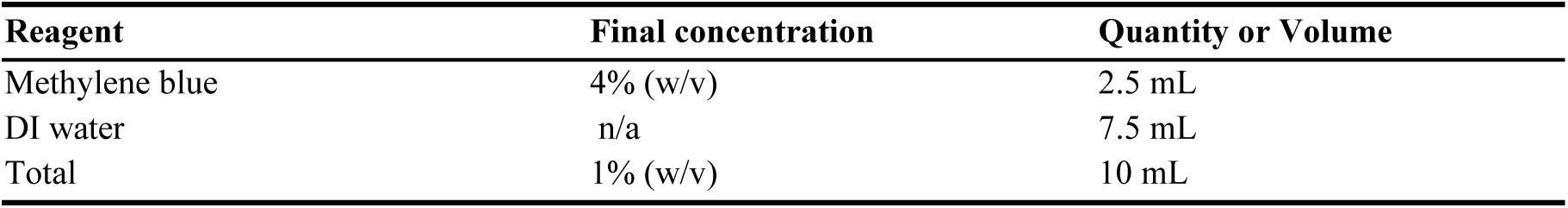
2. **Buffer solution (10 mL) [Used within one week, and stored at 4°C]** 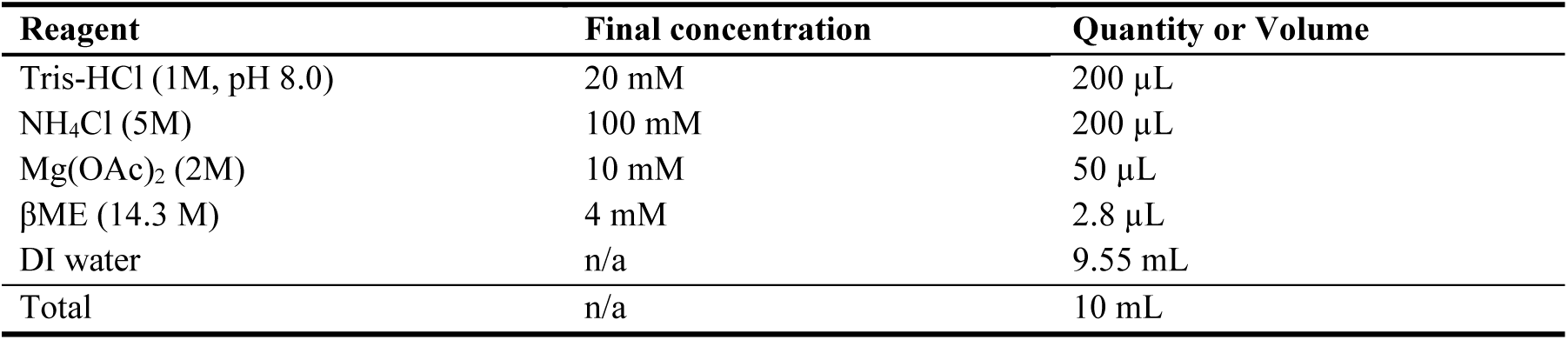

### Laboratory supplies

1. Silicon Wafer with diameter of 100mm (UniversityWafer, Inc. Category: Silicon, ID: 452)
2. Tweezers (Electron Microscopy Sciences, Item number: 0508-L5-PO, https://www.dumonttweezers.com/Tweezer/TweezerStyle/42)
3. AB glue (Loctite, Epoxy, Dries clear, SKU: 1943587, https://www.loctiteproducts.com/products/central-pdp.html/loctite-clear-epoxy/SAP_0201OIL029V3.html)
4. Single-sided tape (Scotch Magic Tape, 0.75 in. x 650 in., https://www.amazon.com/Scotch-Dispensers-Applications-Invisible-Engineered/dp/B0000DH8HQ?ref_=ast_sto_dp)
5. Double-sided tape (Scotch Double Sided Tape, 0.75 in. x 300 in., https://www.amazon.com/Scotch-Double-Dispenser-Standard-237/dp/B0000DH8IT)
6. Aluminum foil (Reynolds, heavy duty, https://www.reynoldsbrands.com/products/aluminum-foil/heavy-duty-foil)
7. Scissors (Fine Science Tools, Surgical Scissors – Sharp-Blunt, Item No. 14001-12, https://www.finescience.com/en-US/Products/Scissors/Standard-Scissors/Surgical-Scissors-Sharp-Blunt/14001-12)
8. PDMS hole punchers (Ted Pella, Inc, Harris Uni-Core, Hole 1.0mm and Hole 0.5mm)
9. Cutting Mat (https://alvindrafting.com/products/hobby-mat-blue-gray, Item No.: HM1218)
10. Glass Cutter Tool Set 2mm-20mm Pencil Style Oil Feed Carbide Tip (Manufacture: MOARMOR, https://www.amazon.com/gp/product/B07Y1D243H/ref=ppx_yo_dt_b_search_asin_title?ie=UTF8&psc=1)
11. Sandpaper (S&F STEAD & FAST: https://www.steadandfast.com/ S&F STEAD & FAST 6 inch Sanding Discs Hook and Loop 24 Pcs, SKU:SA-SC-005)
12. EM grids (TED PELLA, INC., Quantifoil R 0.6/1 holey carbon copper grid, Product No.: 659-300-CU)
13. Tubings (Polymicro Technologies, TSP180350 with Outer diameter (O.D.) 360 µm and inner diameter (I.D.) 180 µm; TSP100170 with O.D. 170 µm and I.D. 100 µm; TSP075150 with O.D. 150 µm and I.D. 75 µm)
14. Pipette tips (1. BRANDTECH, 0.5 – 20 µl, colorless, Category No.: 732104, https://shop.brandtech.com/en/pipette-tips-0-5-20-l-pp-colorless.html; 2. BIOTIX, 20 μL Racked, Sterilized, Part# 63300005 https://biotix.com/products/pipette-tips/xtip4-lts-compatible-pipette-tips/20-%CE%BCl-racked-sterilized-2/)
15. Pipette (Eppendorf, 0.1-5 µL)
16. Ethane gas (Airgas, grade level: research purity)
17. Liquid nitrogen (Airgas, gas grade: Industrial)
18. Vitrification dewar (Nanosoft, SKU: 21021005, https://www.nanosoftmaterials.com/product-page/vitrobot-dewar)
19. Tubes (Thermo Scientific, 15 mL Conical Sterile Polypropylene Centrifuge Tubes, Model No.: 339651, https://www.thermofisher.com/order/catalog/product/339651)
20. Blades (https://www.garveyproducts.com/product/7097, Model No.: CUT-40475)
21. Photo mask (Fineline Imaging, Inc. https://www.fineline-imaging.com/)
22. 500g weight (AMERICAN WEIGH SCALES, model No.: B00SSK3YNO)
23. A polyimide-coated, fused silica capillary tubing (Polymicro Technologies, TSP075150 with O.D. 150 µm and I.D. 75 µm.)
24. SU-8 2050 (KAYAKU ADVANCED MATERIALS INC. SU-8 2050 500ML glass bottle with a cap, Catalog No.: NC0060520, https://www.fishersci.com/shop/products/NC0060520/NC0060520)
25. SU-8 developer (KAYAKU ADVANCED MATERIALS INC. Photoresist developer solution, Catalog No.: NC9901158, https://www.fishersci.com/shop/products/su-8-developer-4l/NC9901158)
26. Polydimethylsiloxane (PDMS; Dow Corning, Sylgard 184, Manufacturer SKU: 4019862)
27. Acetone (Pharmco-Aaper, Midland Scientific, catalog number: 329000000CSGF)
28. Isopropanol (Fisher Chemical, catalog number: BP26184)
29. Acid piranha (a 3:1 mixture of concentrated sulfuric acid (H_2_SO_4_) with hydrogen peroxide (H_2_O_2_), special protection equipment is required for preparing acid piranha[13], and a formal training is highly recommended and required for the sake of safety.)
30. Nitrogen Spray Guns with 0.80 Filter (Cleanroom World, Product Code:TA-NITRO-4-FT, https://cleanroomworld.com/cleanroom-equipment/nitrogen-spray-guns/90-degree-angle-gun-with-hose-assembly-half-inch-tubing) The nitrogen is house-generated in Columbia clean room, the grade level is industrial.
31. Polyimide Tape (MYJOR, Model number: B07RZYY2T1, https://www.amazon.com/MYJOR-Temperature-Protect-Printer-Professionals/dp/B07RZYY2T1?th=1)
32. PEEK tubing (INDEX HEALTH & SCIENCE, Yellow, 1/16” x 0.007” x 5 ft, Part no.: 1536, https://www.idex-hs.com/store/product-detail/peek_tubing_yellow_1_16_od_x_007_id_x_5ft/1536)

## Equipment

1. Optical microscope 1 (Leica, model: LEITZ DM IL) with a digital camera (AmScope, model number: MU 1803-HS)
2. Optical microscope 2 (Nikon, model: ECLIPSE ME600L)
3. Optical microscope 3 (Nikon, model: ECLIPSE LV100ND) with a digital camera (Nikon, model: DS-Ri2)
4. Syringe pump (Cole Palmer Instrument Co., catalog number: 78-0200C)
5. Vacuum system (ZENY™ 3.5CFM 1/4HP Pump: https://www.zeny.us/collections/air-vacuum-pump/, Model: VP 125+; Stainless Steel SlickVacSeal Vacuum Chamber: slickvacseal.com, Brand: SlickVacSeal^TM^)
6. Glow-discharge machine (Ted Pella, Inc. Model: 91000S PELCO easiGlow™ Glow Discharge system for Cryo EM)
7. Plasma processing system for plasma-enhanced chemical vapor deposition (PECVD) (Oxford Instruments Nanotechnology Tools Limited, Model: PlasmaPro®NGP80)
8. Time-resolved apparatus for liquid pumping and pneumatic plunging (developed by Dr. Howard White, Eastern Virginia Medical School). More details about the apparatus can be seen in Section Procedure Subsection D. Preparation of sample for TRCEM and in Figure 13.
9. Hot plate 1 (VWR International, LLC., catalog number: 97042-634)
10. Hot plate 2 (Electronic Micro Systems Ltd., model: 1000 PRECISION HOT PLATE)
11. Hot plate 3 (Electronic Micro Systems Ltd., model: 1000-1 PRECISION HOT PLATE)
12. Spincoating station (ReynoldsTech Fabricators, Inc., programmable FS5.0 spincoating station)
13. Mask aligner (SÜSS MicroTec, SUSS MA6)
14. Plasma System (ANATECH USA https://anatechusa.com/, model: Anatech SCE110)
15. Ultrasonic Cleaner (Branson Ultrasonics Corp., Model No.: B1510R-DTH)

## Software and datasets

1. Nikon NIS-Elements (version 5.21.03, together with Microscope 2, purchased by Columbia University clean room)
2. Fiji/ImageJ (version 2.1.0/1.53c or another version, free and open source)
3. L-Edit (Win32 8.30, license is required), L-Edit is a mask layout editor for Windows-based platforms. For more details about the software, please contact the SIEMENS company: https://eda.sw.siemens.com/en-US/ic/ic-custom/ams/l-edit-ic/.
4. The Electron Microscopy Data Bank (EMDB) (https://www.ebi.ac.uk/emdb/) EMD-29681, EMD-29688, EMD-29687, and EMD-29689

## Procedure

### A. Fabrication of SAR micromixer

Our micromixer is based on the 3-D splitting and recombination (SAR) principle. For 3-D models of the SAR micromixer and its mixing performance, please see the previous work by Feng et al. [14,15]. The fabrication of this PDMS-based micromixer is detailed in the following:

1. Mask design For the fabrication of the micromixer, the conventional soft lithography method is employed. Two layers of microchannel structure (Figure 1A) are required to form a 3-D micromixer, as illustrated by the geometrical model in Figure 1B. Based on the design of micromixer and numerical calculations [11,14], we designed the mask shown in Figures 1C and 1D using the software L-Edit. The photo mask we designed (Figure 1C) has 62 units and they all share the same channel structure as the one shown in Figure 1D, so after the soft lithographic process, we are able to obtain 62 PDMS copies of the same channel structure. When we rotate one layer to make its channel facing the channel of the other layer, we have the two layers shown in Figure 1A.
2. Printing of Photo Mask The mask (Figure 2A) was printed in high resolution (10um @ 32K DPI) by the company Fineline Imaging, a printing company offering the highest-quality laser photoplot films and film photomasks on the market. Its website is: https://www.fineline-imaging.com/. By attaching the photomask film to the glass slide (Figure 2B), we have the mask ready for the following UV lithography. (Note that <u>the Supplementary Mask</u> we designed can be seen in the supplementary information)
3. Fabrication of the microchannels on PDMS material (Note: this process should be conducted in the Yellow Room which is illuminated with yellow light, causing the room to appear yellow and the lamp is designed to filter out harmful UV wavelengths so unwanted exposure of light-sensitive materials such as photoresist is prevented.)
  a. Substrate Preparation: the substrate wafer should be clean and dry to achieve good process reliability. For best results, the silicon wafer is immersed in a piranha solution for 20 mins, followed by a de-ionized (DI) water rinse.
  b. SU-8 coating: 1.) Dispense 5ml of SU-8 photoresist on the wafer and slowly spread it on the wafer by manually tilting the wafer; 2.) Spin at 500 rpm for 10 seconds with acceleration of 100 rpm/second; 3.) Spin at 4000 rpm for 30 seconds with acceleration of 300 rpm/second to achieve a 40µm-thick SU-8 film coating on the wafer. Spincoating station as listed in Equipment section is used in this step.
  c. Soft Bake: put SU-8 coated wafer on the hotplate and heat up first to 65°C for 3 min and then up to 95°C for 6 min, then cool down to room temperature.
  d. Exposure: use the photomask (Figure 2C) and apply exposure dose of 160 mJ/cm^2^ using Mask aligner (SÜSS MicroTec, SUSS MA6).
  e. Post-Exposure Bake: put the wafer on the hotplate and heat up to 65°C for 1 min, then up to 95°C for 6 min, then cool down to room temperature.
  f. Development: spray KAYAKU SU-8 developer onto the SU-8 wafer within 5 min, then clean and clear structures coming up.
  g. Rinse and Dry: spray and wash the developed image with fresh solution of Isopropyl Alcohol (IPA) for 10 seconds. Air-dry in a gentle way with nitrogen gas.
  h. Hard-Bake (final cure): bake temperature 160°C for 10 minutes. The SU-8 mold is obtained as shown in Figure 3A.
  i. Use aluminum foil to form a container with the SU-8 mold on the bottom, use the polyimide tape to surround the edge of the wafer and to fix it in place (Figure 3B).
  j. 30g base and 3g curing agent are mixed in a cup and the mixture (∼30.5g) is poured into the container mentioned above.
  h. Put the container into the vacuum for degassing and cure it on the hotplate at 100°C for 1 hour and then cool down the PDMS.
  i. Peel off the cured PDMS replica from the SU-8 mold as shown in Figure 3C. (Note: the SU-8 mold can be reused for a couple of years if it is properly preserved)
  j. Follow the boundary of each PDMS unit, as highlighted in Figure 3C and cut the PDMS replica into pairs of PDMS slabs with the same microchannel structure as shown in Figure 1D, and each pair (Figure 3D) is ready for the micromixer fabrication. (Note: one PDMS replica (Figure 3C) from the SU-8 mold produces 31 pairs of PDMS slabs, which ideally enables us to obtain 31 micromixers.)
4. Alignment and bonding of Micromixing channel
  a. Clean the surface of the PDMS with tape.
  b. Plasma-treat the surfaces of two PDMS slabs with the microchannel side up (15s Ar at 100W and 1min O_2_ at 100W).
  c. Apply a drop of Deionized (DI) water (4µL) on the surface of the bottom layer for lubrication.
  d. Align the microchannels on the PDMS slabs under the microscope as shown in Figure 4. (Note that a lot of practicing is required, and the alignment should be completed within 1 min for best bonding quality).
  e. Put the aligned PDMS slabs in vacuum (<50 mtorr) for 2 mins after alignment to remove extra water in between the PDMS slabs; this procedure pre-bonds the PDMS slabs.
  f. Check if misalignment has occurred as a result of the exposure to the vacuum.
  g. Cover the pre-bonded PDMS slabs with small pieces of aluminum foil (Figure 5A).
  h. Bake pre-bonded PDMS slabs on the hotplate at 120°C for 10 min with a 500g weight on it (Figure 5B) to further strengthen the bonding, and then check the alignment under the microscope. This is the fully formed micromixer.
  i. Apply plasma treatment to the surfaces of small glass slide and the micromixer (15s Ar at 100W and 1min O_2_ 100W).
  j. Apply a small drop of DI water (2µL) on the surface of the glass slide for lubrication.
  k. Put the bonded PDMS slab onto the surface of the glass slide and move it to its center.
  l. Bake at 120°C for 2 min.
  m. Cut the tubing (O.D.: 150 µm and I.D.:75 µm) into pieces (Figure 5C) and sand the openings using sandpapers (3000 grit and then 4000 grit) such that the ends will become smooth and flat (Figure 5D).
  n. Insert well-polished tubings (Figure 5D) into the two inlets and the outlet of the micromixer (Figure 5E). Here a polyimide-coated, fused silica capillary tubing (O.D. 150 µm; I.D. 75 µm) is used. Note that another type of tubing O.D. 170 µm; I.D. 100 µm also can be used for this step, and the length of the tubing depends on the time point desired. Note that the tubings with 2.5 mm, 4.5 mm, and 5 mm in length are used according to our need.
  o. Apply a very tiny amount of AB glue (size of bead: ∼ 1 mm in diameter) on the areas between the tubing and the micromixer. The area is marked by the dashed curve (Figure 5E-F). The glue is filled between the tubing and the PDMS channel, as shown in Figure 5H-J. (Note that after mixing the AB glue, wait for 1 min before applying it to the target area to prevent the glue from flowing too quickly and entering the tubing, thereby blocking it).
5. SiO_2_ coating
  a. Put the PDMS micromixers (as shown in Figures 5C and 6A) into the chamber of the PlasmaPro®NGP80 PECVD machine.
  b. Stabilize the stage temperature at 300°C.
  c. Let in the source gasses SiH_4_ (170 sccm) + N_2_O (710 sccm) at a vacuum pressure of 200 mTorr.
  d. Apply high radio frequency (RF) power (50W) to create plasma inside the process chamber. Set the coating strike time for 20 min.
  e. Vent and open the chamber, and store the coated micromixers in petri dish in normal room for future use.
6. Testing of the micromixer
  a. Use the IDEX PEEK tubing of 6 mm in length as connector to connect one end of the polyimide tubings (O.D. 150 µm; I.D. 75 µm, 30 cm in length) to inlet of the micromixer (Figure 7A) and the other end is inserted into another piece of the IDEX PEEK tubing of 4 cm in length, which will be used to be connected to the syringes pump, and apply the AB glue on the junction regions – the process is similar to the step as shown in Figure 5E.
  b. Build the setup for testing, which includes a syringe pump and a microscope with a camera and laptop (Figure 7B).
  c. Connect the micromixer inlets to the syringes via glass tubings with O.D. 150 µm and I.D. 75 µm.
  d. Connect the micromixer outlet to a tube that collects the mixture solution.
  e. Introduce the dye solution (or fluorescent solution) and DI water into the micromixer.
  f. Run the total flow rates at 2, 4, 6, 8, 9 µL/s (Figure 8B-F) and capture mixing status for each setting.
  g. Check if there is evidence of any expansion in the mixing area of microchannels at 9 µL/s (540 µL/min), which is 50% higher than the intended working flow rate (6 µL/s).
  h. Check if there is obvious leakage at 9 µL/s.
  i. Measure the intensity distribution across the outlet area (marked by dash lines in Figure 8B-F). As one example, based on Figure Figure 8D, we obtained the intensity distributions at inlet and outlet at a total flowrate of 6 µL/s as shown in Figure 8G. We found that the intensity at the outlet becomes uniformly distributed after mixing. For the calculation of the mixing efficiency, we refer to Feng et al.[14].
  j. Run each flow rate (as shown in Figure 8B-F) five times, each time for four seconds, to check if the micromixer is durable enough.
  k. Use only those micromixers that pass this high flow-rate quality check. Discard the others.

**Figure 1.**
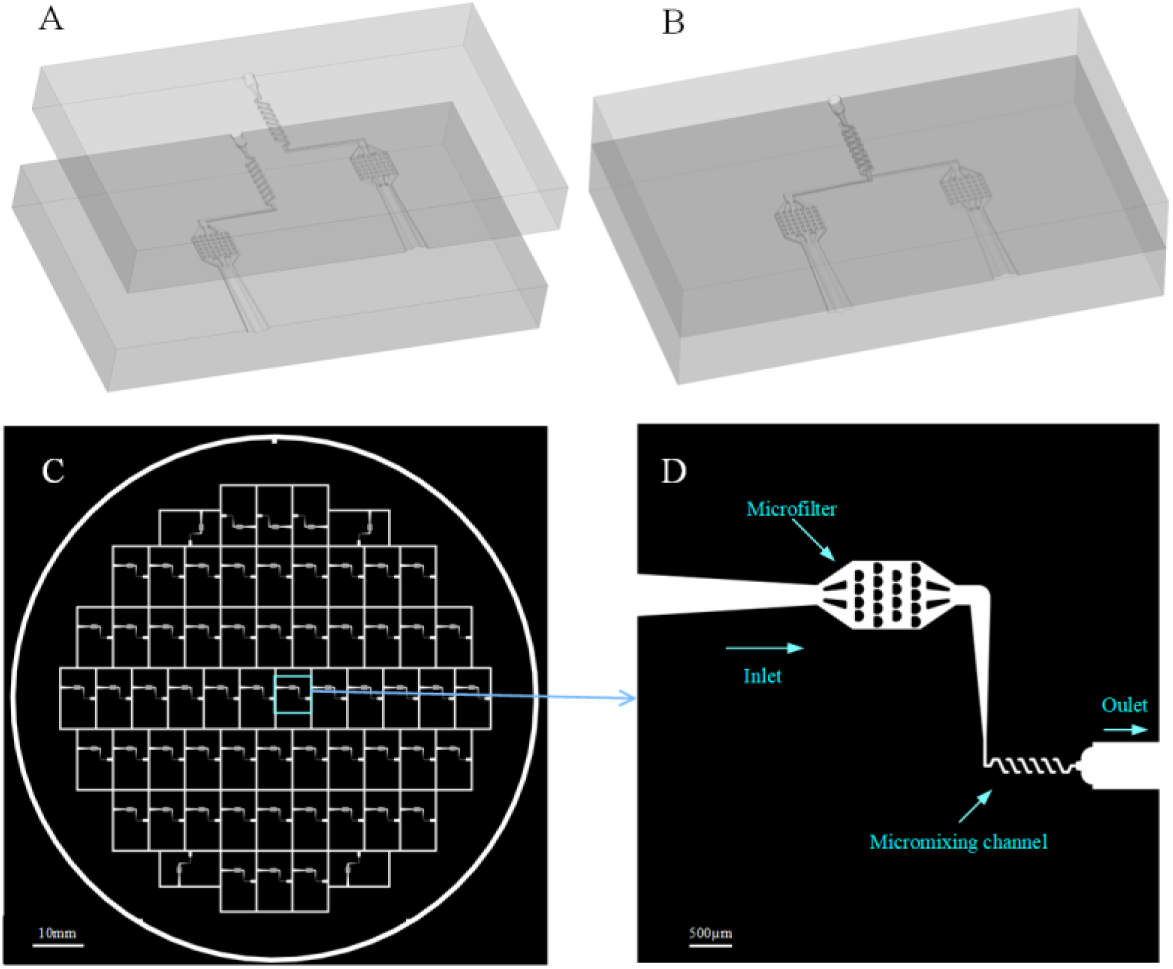
The design of the photo mask. A. Two layers of microchannel structure, which are made by the same photoresist and are thus identical. B. Geometrical model of the 3-D micromixer. C. A photo mask with 62 units. D. One unit of the mask (C) zoomed in, where the inlet with a microfilter was designed to capture big particles.

**Figure 2.**
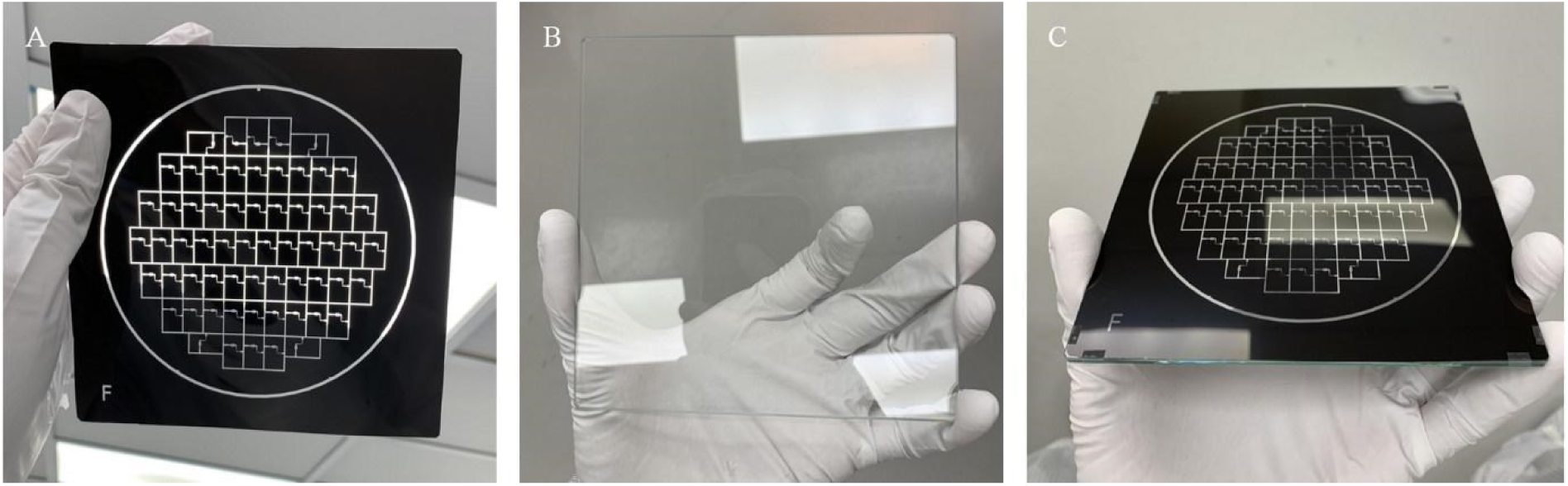
Photo mask printed by high-resolution printer from Fineline Imaging. A. Photo mask. B. Glass slide. C. The photo mask attached to the glass slide.

**Figure 3.**
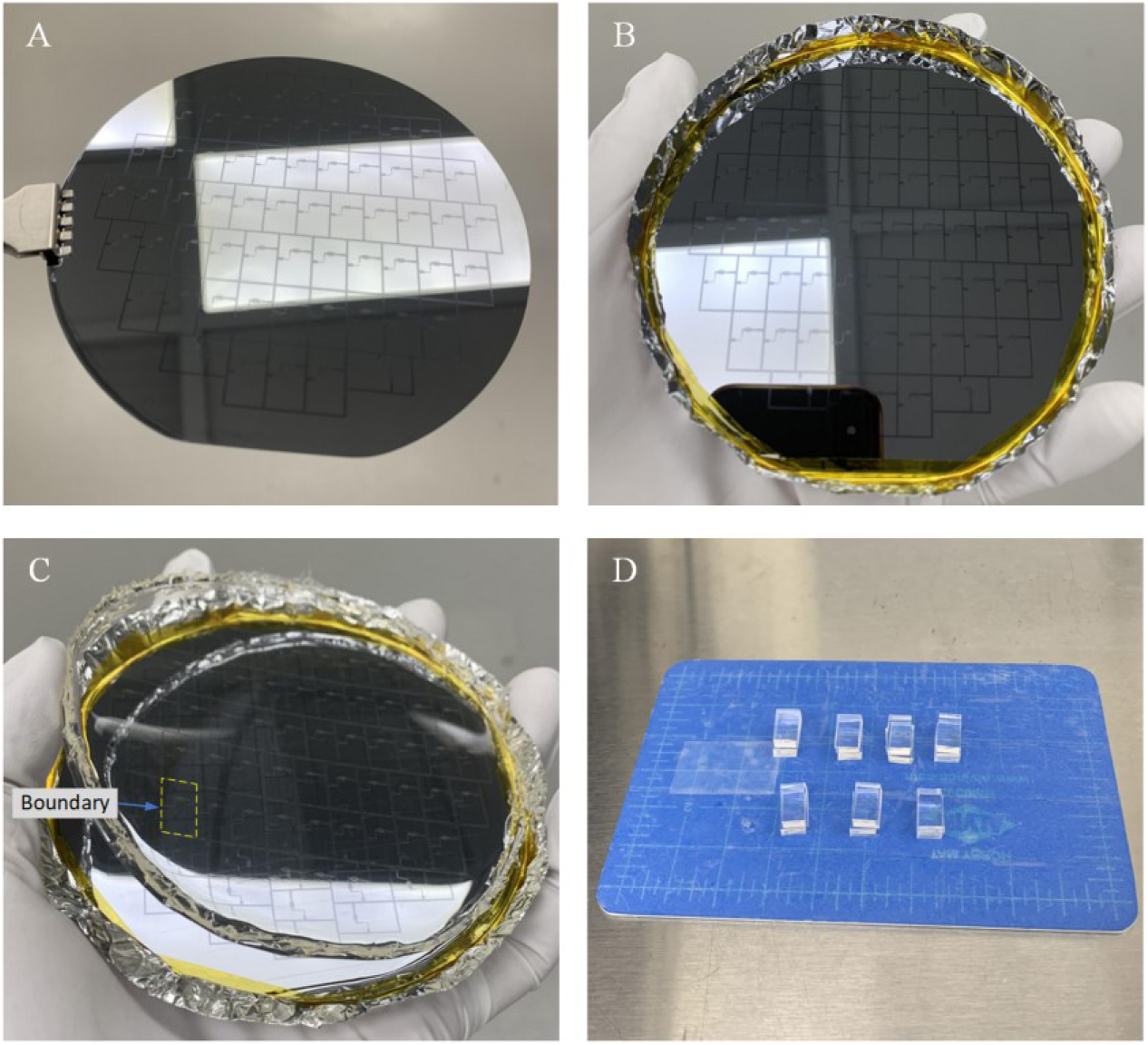
Fabrication of the microchannels from PDMS material. A. SU-8 mold. B. Container with SU-8 mold on the bottom. C. PDMS replica peeled off from the SU-8 mold, with the boundary of a PDMS unit highlighted. D. Example of seven pairs of PDMS slabs. (Note that the white areas in A, B and C are from the reflection of the ceiling light, not from patterns on the mold).

**Figure 4.**
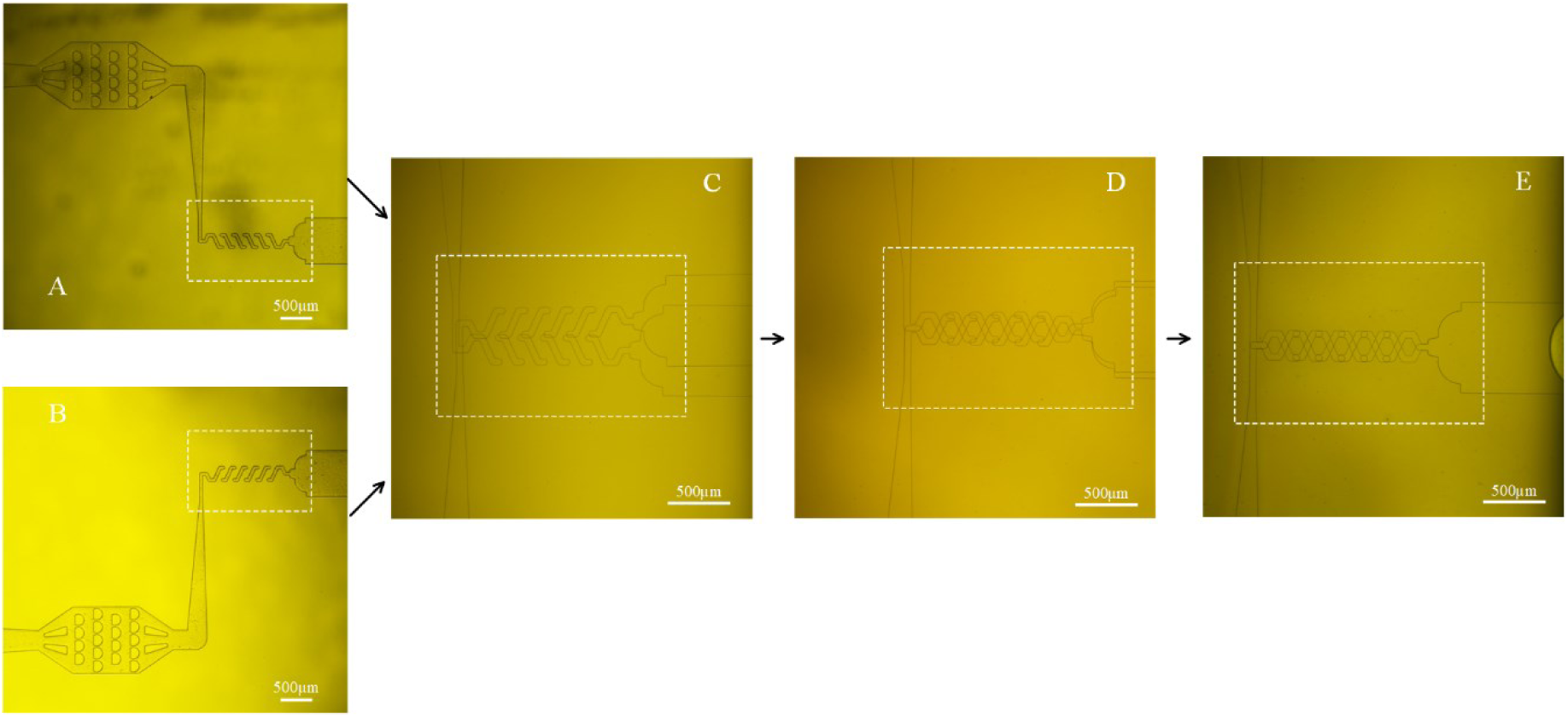
Alignment during the fabrication of the 3-D PDMS micromixer. A, B. Top and bottom layer of microchannel. C. Misalignment in the beginning. D. Move top layer to align the microchannel with reference to inlets and outlet. E. Well-aligned microchannel.

**Figure 5.**
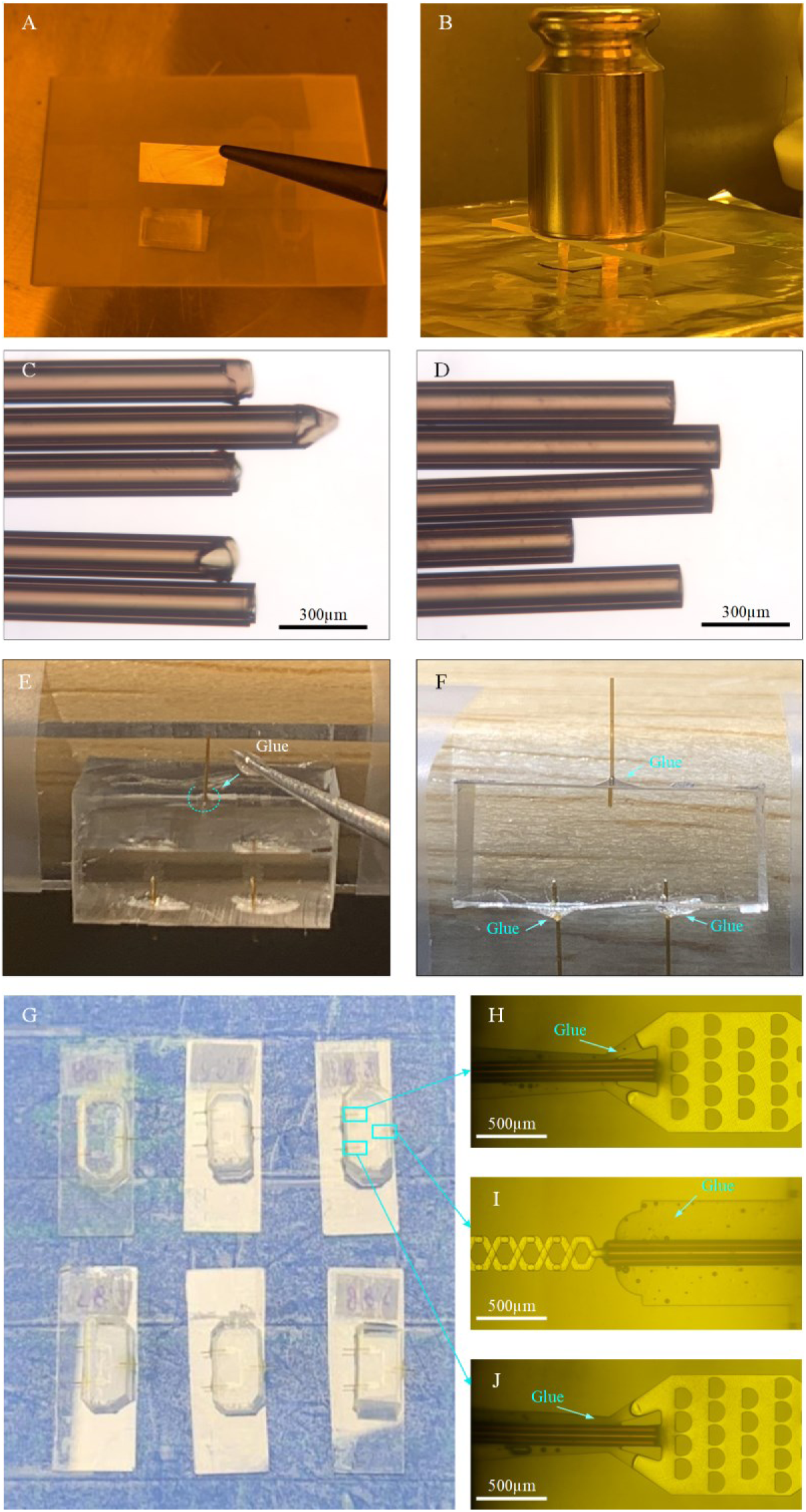
Fabrication of the micromixer. A. Cover the aligned PDMS slabs with small pieces of aluminum foil. B. Put a 500g weight on the aligned PDMS slabs during baking. C. Raw tubing (O.D.: 150 µm and I.D.: 75 µm) with coarse opening. D. Sanded tubing (O.D.: 150 µm and I.D.: 75 µm); the length depends on the need. E. Application of a bead of glue onto the corner between the tubing and PDMS surface, as marked by the dashed curve. F. The glue covers the corner after application. G. Examples of six micromixers with inlet and outlet tubings inserted and glued. H, J. Two tubings are inserted and connected to the microfilters and well glued. I. One glass capillary tubing is inserted, connected to the outlet and well glued.

**Figure 6.**
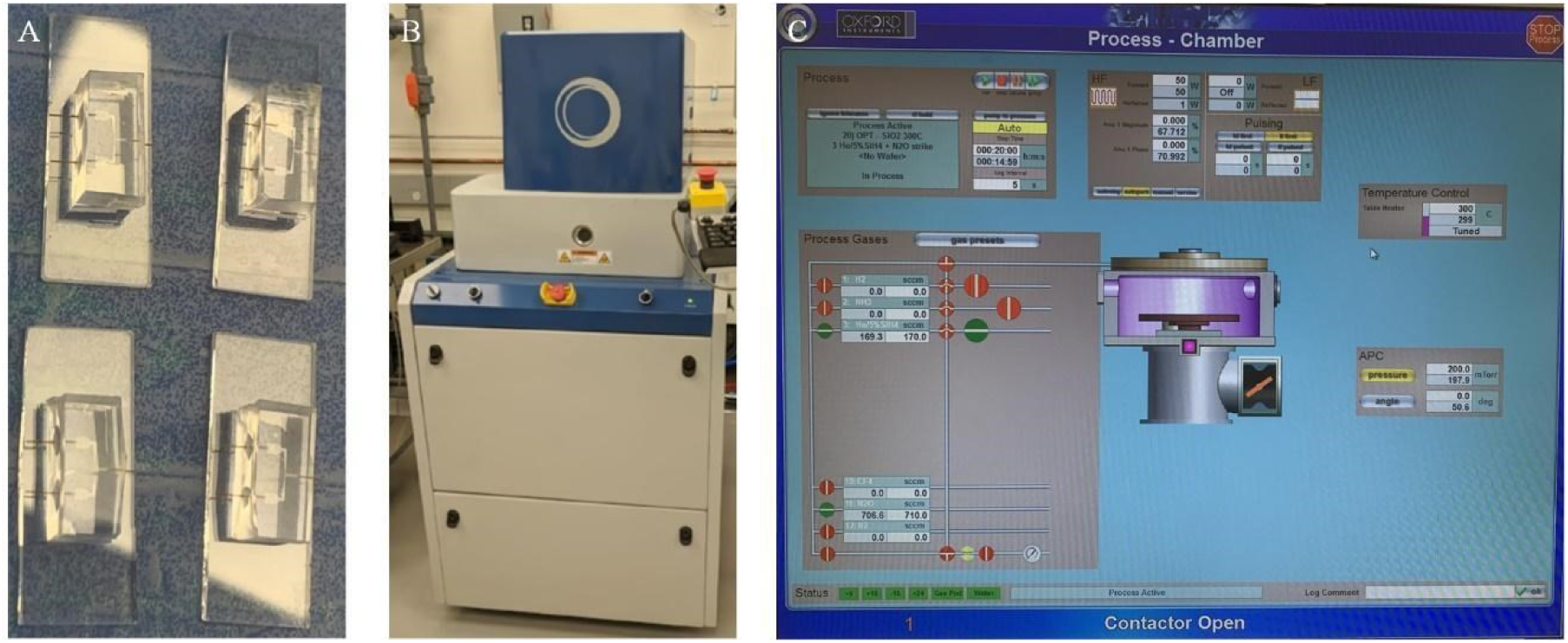
**SiO**_2_ **coating.** A. Examples of four micromixers fabricated following above-mentioned step 4. B. The PECVD machine of model PlasmaPro®NGP80. C. Monitoring of the coating process.

**Figure 7.**
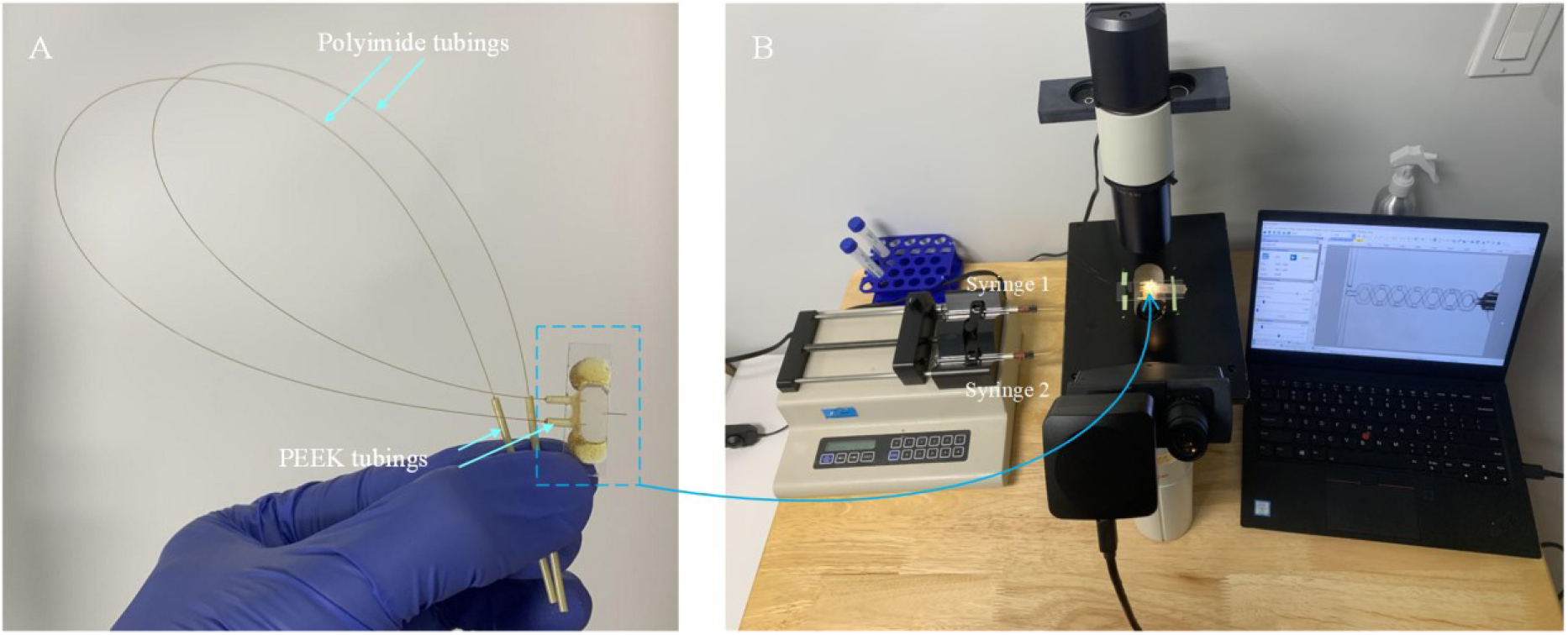
Setup for testing the micromixer. A. Fabricated micromixer with tubings for connection to the syringe pump. B. Equipment required for testing, comprising of a syringe pump, a microscope, a camera and a laptop.

**Figure 8.**
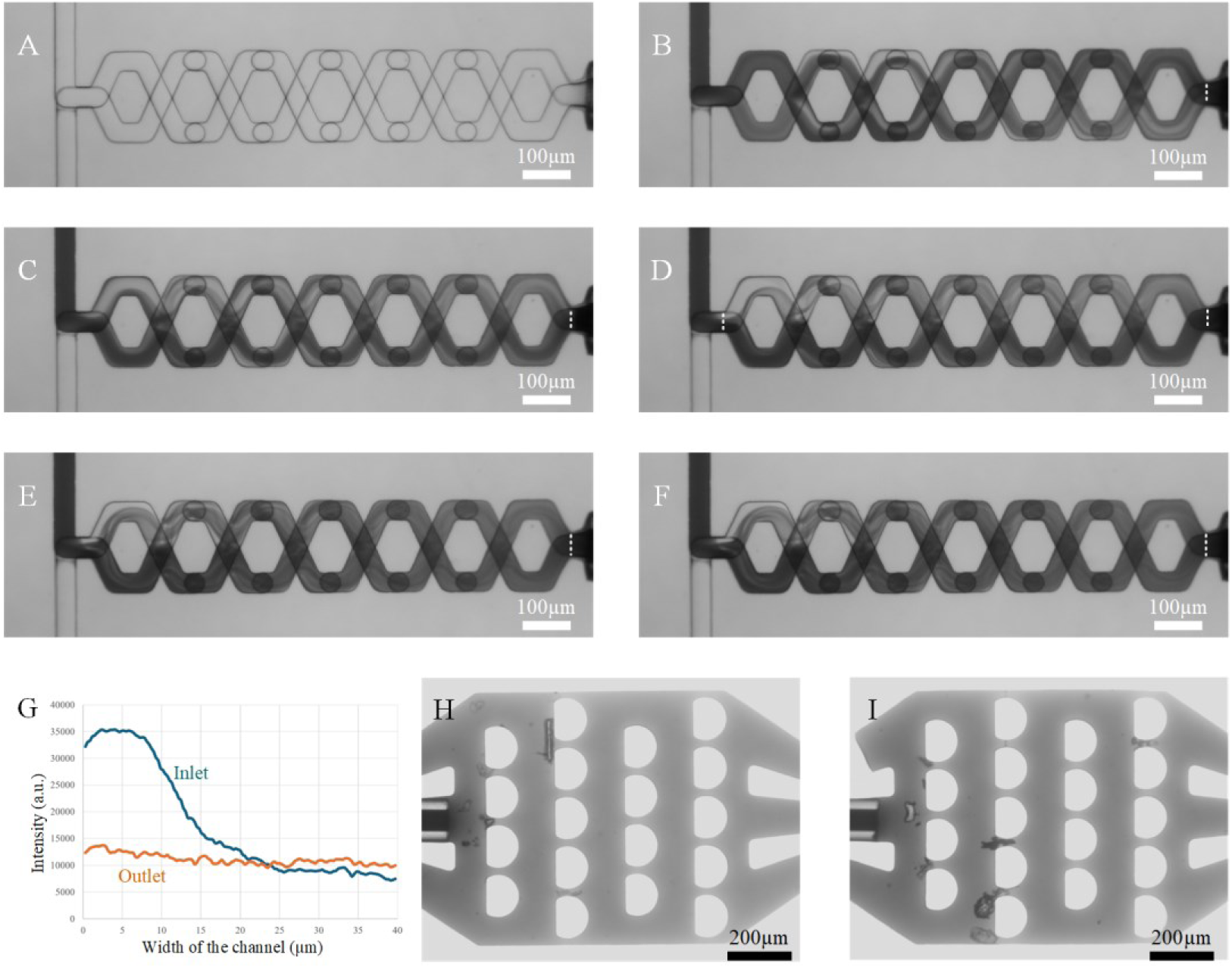
Testing of micromixer. A. The micromixing channel at flowrate of 0 µL/s. B, C, D, E, and F images of mixing statuses at flowrates of 2, 4, 6, 8, and 9 µL/s, respectively. G. Intensity distributions at inlet and outlet when the flowrate is 6 µL/s as shown in Figure 8D. H, I. Microfilters at the inlets, for capturing big particles above 20 µm in diameter.

### B. Fabrication of microsprayer

1. Cut the PDMS into small pieces of PDMS slab (length x width x thickness: 6 x 5 x 4 mm) and use PDMS punchers to make holes to accommodate sprayer outer tube, liquid tube and gas tubes (Figure 9A). (Note: 1–the PDMS slab used here is plain, without any microfluidic channel. 2–some practicing will be required for cutting PDMS and punching holes with PDMS hole punchers of 1 mm and 0.5 mm in diameter, respectively.)
2. Cut the normal glass slide (length x width x thickness: 75 x 25 x 1 mm) into small pieces with length x width x thickness: 25 x 7 x 1 mm (Figure 9B) using a glass cutting tool. (Note: different glass cutters tools can be used, and some practicing will be required)
3. Apply plasma treatment to the surface of PDMS slab (Figure 9A) and small glass slide (Figure 9B) (2min, 15mA at air in EasiGlow cleaning machine).
4. Apply a small drop of DI water (2µL) on the surface of small glass slide for lubrication. Align the PDMS slab on the glass slide as illustrated in Figure 9C and then bake the assembly on hot plate at 100°C for 2 min for strong bonding.
5. Cut the BRANDTECH pipette tips into a shape as shown in Figure 9D and insert them into the side holes of the PDMS slab. They will serve as gas inlets for the microsprayer. After plasma treatment on it (2min 15mA at air in EasiGlow cleaning machine), apply some glue (size of bead: ∼ 5 mm in diameter) on the junction area of the connection (Figure 9E) and wait for the glue to dry.
6. Cut the tubing (O.D.: 350 µm and I.D.: 180 µm) into pieces with length of 6 mm (Figure 9F) and sand the openings using sandpapers (3000 grit and then 4000 grit) such that the ends will become smooth and flat (Figure 9G). In the following we refer to the tubing obtained as “tubing 1”.
7. Insert tubing 1 into the hole of the PDMS slab as shown in Figure 9H. Then apply a tiny amount (approx bead size: ∼ 1 mm in diameter) of glue on the junction area between the PDMS and tubing 1 and wait for the glue to dry. (Note: there is one raw tubing (O.D.: 170 µm and I.D.: 100 µm) inside tubing 1, which is used to check if tubing 1 is roughly concentric with the hole. It will be removed later.)
8. Cut the tubing (O.D.: 170 µm and I.D.: 100 µm) into pieces with length of 10 mm (Figure 9I) and sand the openings using sandpapers (3000 grit and then 4000 grit) such that the ends will become smooth and flat (Figure 9J). In the following we refer to the tubing obtained as “tubing 2.”
9. Insert tubing 2 into tubing 1 and move tubing 2 until 1 mm of it protrudes outside PDMS on nozzle side, then apply a tiny amount (size of bead: ∼ 3 mm in diameter) of glue on the junction area between tubing 1 and tubing 2 (Figure 9K), and wait for the glue to dry. Figure 9L shows several copies obtained by repeating the steps above.
10. Cut the BRANDTECH pipette tips into a shape with length of 2mm as shown in Figure 9M, to be used as the microsprayer outer tube. In the following we refer to the tubing obtained as “tubing 3.”
11. Insert tubing 3 into the hole of PDMS slab (Figure 9N) and carefully align tubing 3 with tubing 2 under the microscope to make them concentric, as shown in Figure 9O. (Note: tubings 2 and 3 form the microsprayer, with tubing 2 for the liquid and tubing 3 for the driving gas.)
12. Follow step 11, plasma treatment on sprayer assembly (2min 15mA at air in EasiGlow cleaning machine), apply some glue (bead size: 1mm in diameter) to the junction area between tubing 3 and rim of PDMS hole and wait for the glue to dry. Figure 9P shows several copies obtained by repeating the steps above.
13. Cut the soft tubing (O.D.: 2.4 mm and I.D.: 0.8 mm) into pieces with length of 15 cm. And cut the BRANDTECH pipette tip with a length of 8mm and bend them to be in a curved shape. Assemble the soft tubing and the curved tip as shown in Figure 9Q. In the following we refer to the tubing obtained as “gas tubing.”
14. Insert the two gas tubings into the two gas inlets of the microsprayer (Figure 9R) and apply some glue (bead size: ∼ 4 mm in diameter) to the junction area between gas tubings and gas inlets, and wait for the glue to dry.
15. Now the microsprayer is ready to be used in the fabrication of the entire chip assembly with time point longer than 20ms.

### C. Fabrication of the microfluidic chip assembly

#### C1: For time points longer than 20 ms

1. Choose proper glass capillary tubing to connect the micromixer with the microsprayer. Here for demonstration a tubing with O.D. 350 µm and I.D. 180 µm is used and cut to a length of 83 mm, and we will call it the microreactor tubing. (Note: since the reaction time is related to the volume of the microreactor tubing, the length and diameter of the microcapillary tubing can be changed to achieve different time points. The micromixer outlet tubing and microsprayer inlet tubing must be changed in size for proper connection.)
2. Insert the outlet tubing of the micromixer (O.D. 170 and I.D. 100 µm) into the microreactor tubing (O.D. 350 µm and I.D. 180 µm) on one end and insert liquid tubing of the microsprayer into the microreactor tubing (O.D. 350 µm and I.D. 180 µm) on the other end. Thus the micromixer, the microreactor, and the microsprayer are now connected to each other into the complete assembly (Figure 10).
3. Apply a tiny amount (bead size: ∼ 1 mm in diameter) of glue to the connection area between the micromixer and the microreactor, and the area between the microreactor and microsprayer, and then wait for the glue to dry. (Note: after mixing the AB glue, wait for 1 min to reduce the fluidity of the glue before applying it to the areas of interest, to prevent the glue from clogging the microreactor tubing. Regarding the timing for applying the glue, first do some testing on separate raw tubings with the same diameters as specified in steps 1 and 2 is required to check how the glue flows inside the gap between the tubings and when it stop s flowing).
4. Estimate and calculate the reaction time based on the design of the fabricated chip. As we stated in our work[11], the total reaction time is comprised of the mixing time in the micromixer, the reaction time in the micro-capillary reactor, the spraying time, the plunging time, and the reaction time during the vitrification process. In this example (Figure 10), a total reaction time of 350 ms will be obtained. (Notes: 1–the volume taken up by the inserted tubing on the connection region needs to be subtracted. 2–In our work[11], following this process, we fabricated a group of chips with three time points: 25 ms, 140 ms, and 900 ms.).
5. Check the sample adsorption with the *E. coli* 70S ribosome by using the setup as depicted in Figure 13. Collect the sample before and after passing the device, and measure the concentration by using our Nanodrop UV-Vis Spectrophotometer. In this testing, 94% of the initial concentration can be retained using the SiO_2_-coated chip. (Gas pressure for the spray is 8 psi and the total liquid flowrate is 6 µL/s; for more details on different coatings see ref[11]).

#### C2: For time points shorter than 20 ms

1. Follow the step shown in Figure 9A to prepare the PDMS slab with holes (one front hole, two side holes and one back hole), and trim it to have a smaller size: length x width x thickness: 5 x 4 x 2 mm. Here we call this PDMS slab a nozzle PDMS slab. The front hole is used for accommodating the sprayer outer tube, the back hole is for the liquid tube, and the side holes are for gas tubes (Figure 11A).
2. Cut the BRANDTECH pipette tip with a length of 8 mm and bend them into a curved shape. Then insert the tip into each side hole of the nozzle PDMS slab as shown in Figure 11B.
3. Take one micromixer with a liquid tube as its outlet (Figure 5G), and insert the liquid tube into back hole of the nozzle PDMS slab, then the two surfaces of the nozzle PDMS and micromixer PDMS slab can make contact with each other as shown in Figure 11C.
4. Apply plasma treatment to assembly of micromixer and nozzle PDMS slab (Figure 11C) (2 min, 15 mA at air in EasiGlow cleaning machine). Then fix the micromixer on the side of table to ensure that the front hole of nozzle PDMS slab faces up (Figure 11D), and apply some AB glue (bead size: 1 mm in diameter) a few times until the AB glue will cover the corner between the nozzle PDMS and micromixer PDMS slab, as dash-circled in Figure 11E. After the glue becomes dry, the nozzle PDMS is attached to the micromixer (Figure 11F).
5. Cut the BRANDTECH pipette tips into a shape with length of 2.5 mm as shown in Figure 11G, to be used as the microsprayer outer tube.
6. Insert the outer tube into the front hole of the PDMS slab (Figure 11H) and carefully align the outer tube with the liquid tube under the microscope to make them concentric, as shown in Figure 11I.
7. Fix the micromixer on the side of a table to ensure that the outer tube of the microsprayer faces up (Figure 11J), and apply some AB glue (bead size: 1 mm in diameter) a few times until the AB glue will cover the corner between outer tube and the nozzle PDMS slab (Figure 11K). After the glue becomes dry, the outer tube is attached to nozzle PDMS slab and a microsprayer is formed.
8. Cut the IDEX PEEK tubing (O.D. 1/16”, I.D. 0.007”) into pieces with length of 6 mm, which serve as liquid tubing connector, and put two pieces on the inlets of the micro-mixer as shown in Figure 11L.
9. Cut the soft tubing (O.D.: 2.4 mm and I.D.: 0.8 mm) into pieces with length of 15 cm. And cut the BIOTIX pipette tip with a length of 5 mm. Assemble the soft tubing and the tip (as shown in Figure 11M) to serve as gas tubing.
10. Insert the two gas tubings into the two gas inlets of the micro-sprayer (Figure 11N).
11. Use tubings (O.D. 150 µm; I.D. 75 µm) with proper length (30 cm) as liquid tubing, and insert one end into liquid tubing connectors for connection to inlets of the micromixer (Figure 11O).
12. Apply some glue (bead size: ∼ 4 mm in diameter) to the junction area between gas tubings and gas inlets, to the junction area between micro-mixer and liquid tubing connectors, and to the junction area between liquid tubing connectors and liquid tubing, as dash-circled in Figure 11P, and wait for the glue to dry.
13. Insert the other end of the liquid tubing into a piece of the IDEX PEEK tubing of 4 cm in length (Figure 11R), which will be used to be connected to the syringes pump, and apply the AB glue on the junction regions, and the process is similar to the step as shown in Figure 5E.
14. At last we now have the microfluidic chip assembly with time point shorter than 20 ms (Figure 12). (Note that in our work[11], following this process, we fabricated the chip assembly with a time point of 10 ms.)

### D. Estimation of the reaction time obtained using this protocol

Based on our previous study[11], the total reaction time can be estimated as *t*=*t_*m*_*+*t_*r*_*+*t_*f*_*+*t_*p*_*+*t_*v*_*, where *t_*m*_* is the mixing time in the micromixer, *t_*r*_* is the reaction time in the micro-capillary reactor, *t_*f*_* is the flight time in the spray, *t_*p*_* is the plunging time, and *t_*v*_* is the reaction time during the vitrification process. Briefly, each part is described as follows:

1. *t_*m*_* is estimated by the equation: *t_*m*_*=*V_*m*_*/*U_*f*_*, *V_*m*_* is the volume of micromixer, *U_*f*_* is the total flowrate of the solutions introduced into the micro-mixer. Here *V_*m*_* is 2.80 nL for this micromixer, and *U_*f*_* is 6 μL/s, so the mixing time is estimated to be ∼0.47 ms.
2. *t_*r*_* is estimated by the equation: *tr*=*L_*r*_*/<u>*V*</u>, where *L_*r*_* is the length of the microcapillary reactor, <u>*V*</u> is the mean velocity of the fluid.
3. *t_*f*_* is estimated by the equation: *t_*f*_*=*D_*f*_*/*V_*d*_*, *V_*d*_* is the averaged velocity of the droplets, and *D_*f*_* is the distance from the micro-sprayer orifice to the EM grid, fixed at 3.5 mm for our implementation. When liquid flowrate is 6 μL/s and gas pressure is around 8psi, *V_*d*_* is around 6.4 m/s[11], so that we obtain a mean droplet flying time of ∼0.55 ms.
4. *t_*p*_* is estimated by the equation: *t_*p*_*=*H_*p*_*/*V_*p*_*, where *V_*p*_* is the plunging speed and *H_*p*_* is the plunging height from the micro-sprayer orifice to the liquid ethane surface. In this protocol, *V_*p*_*=1.9 m/s at N_2_ gas pressure of 40 psi, and *H_*p*_* is 10 mm or 15 mm. At *H_*p*_* of 10 mm, *t_*p*_* is ∼5.26 ms; at *H_*p*_* of 15 mm, *t_*p*_* is ∼7.89 ms.
5. *t_*v*_* is estimated by the equation: *t_*v*_*=(*T*_*r*oom_−273.15)/CCR, where *T*_*r*oom_ is the room temperature, and CCR is the critical cooling rate, which is ∼10^5^ K/s. So we can estimate the reaction time in the vitrification process, which amounts to ∼0.23 ms.

**Figure 9.**
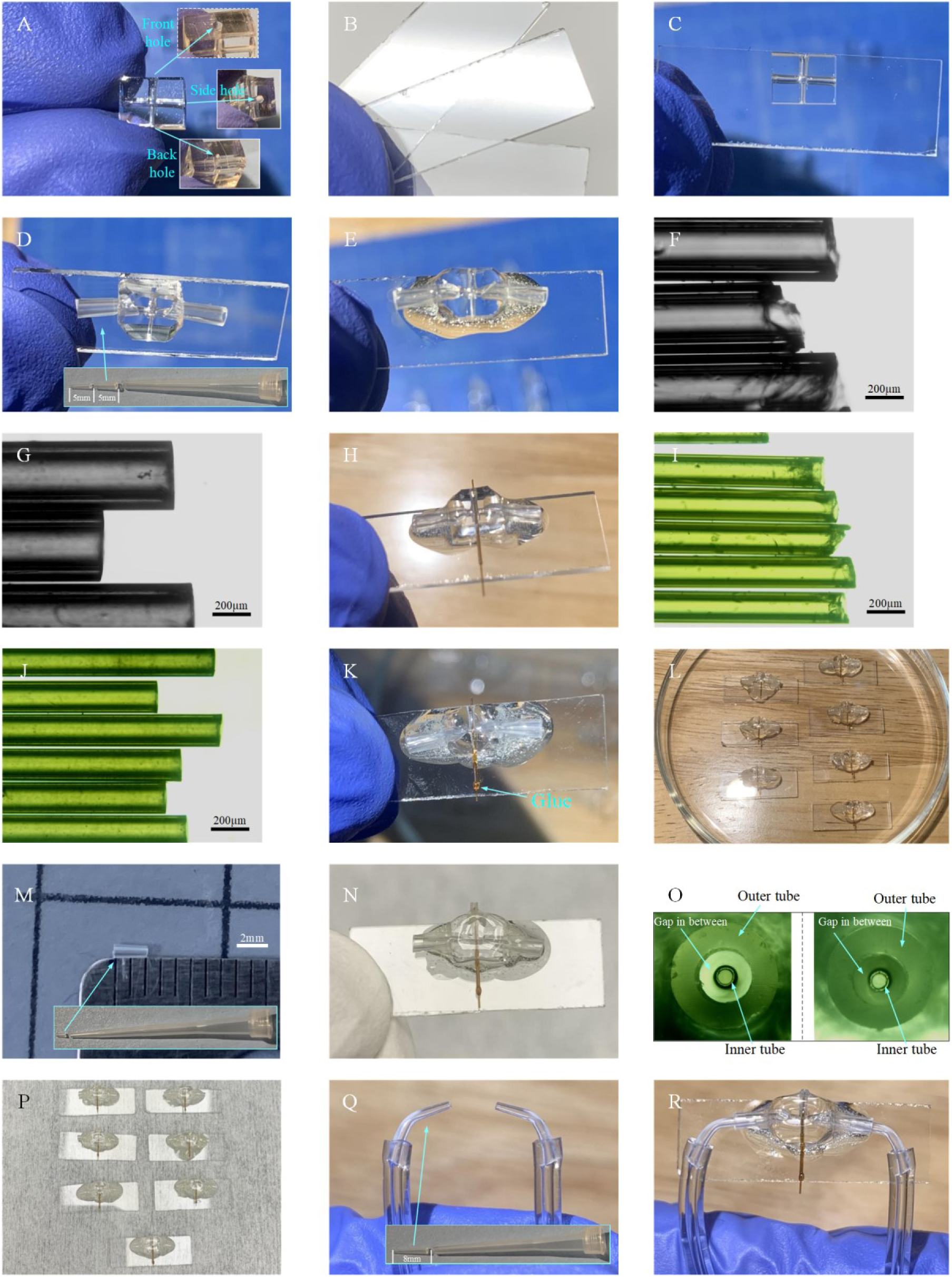
Process of microsprayer fabrication. A. Trimmed PDMS slab with holes for holding the gas inlet tubings, liquid inlet tubing, and micro-sprayer nozzle. B. Small pieces of glass slide. C. Bonding of the PDMS slab on the glass slide. D. Insertion of gas tubings into the PDMS slab. E. Application of glue to seal the connection area between the gas tubings in the PDMS slab. F. Raw tubing (O.D.: 350 µm and I.D.: 180 µm) with coarse opening. G. Sanded tubing (O.D.: 350 µm and I.D.: 180 µm, length: 6 mm, named: tubing 1), which is used for holding the tubing 2. H. Insertion of tubing 1 into the hole of the PDMS slab, with one raw tubing (O.D.: 170 µm and I.D.: 100 µm) inside tubing 1. This procedure is used to check if tubing 1 is roughly concentric to the PDMS hole. I. Raw tubing (O.D.: 170 µm and I.D.: 100 µm) with coarse opening. J. Sanded tubing 2 (O.D.: 170 µm and I.D.: 100 µm, length: 6 mm) serving as the liquid tubing. K. Insertion of tubing 2 into tubing 1. L. Example of seven copies obtained by repeating the steps above. M. Outer tubing 3 of microsprayer with a length of 2 mm. N. Insertion of tubing 3 into the PDMS slab to form the microsprayer nozzle. O. Tubings 3 and 2 are aligned to be concentric and the tubing 2 as the inner tube for the liquid and tubing 3 as the outer tube for the driving gas. P. Example of seven microsprayers obtained by repeating the steps above. Q. Gas tubings. R. Assembled microsprayer ready for use.

**Figure 10.**
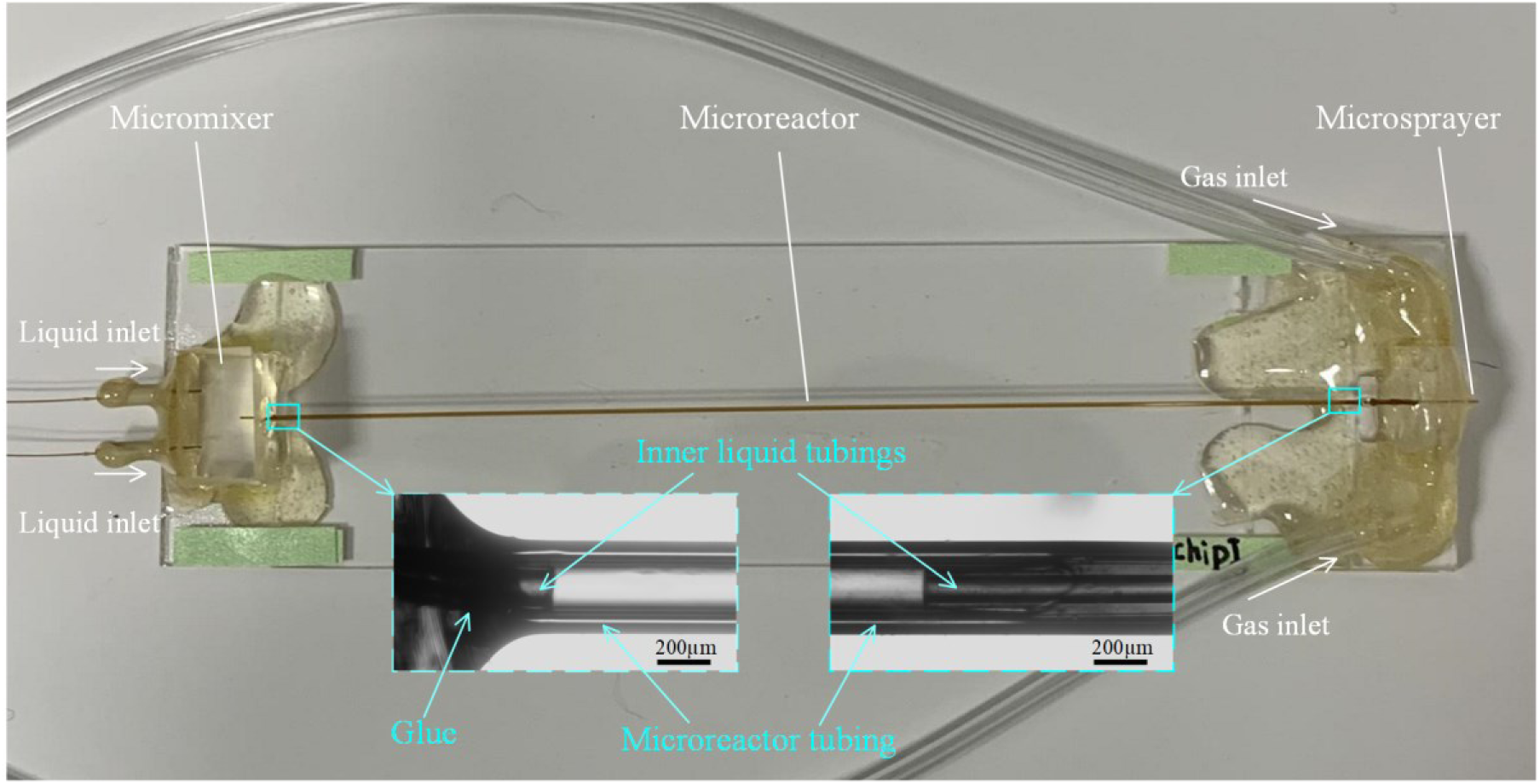
**The fabrication of microfluidic chip assembly with time points longer than 20ms.**

**Figure 11.**
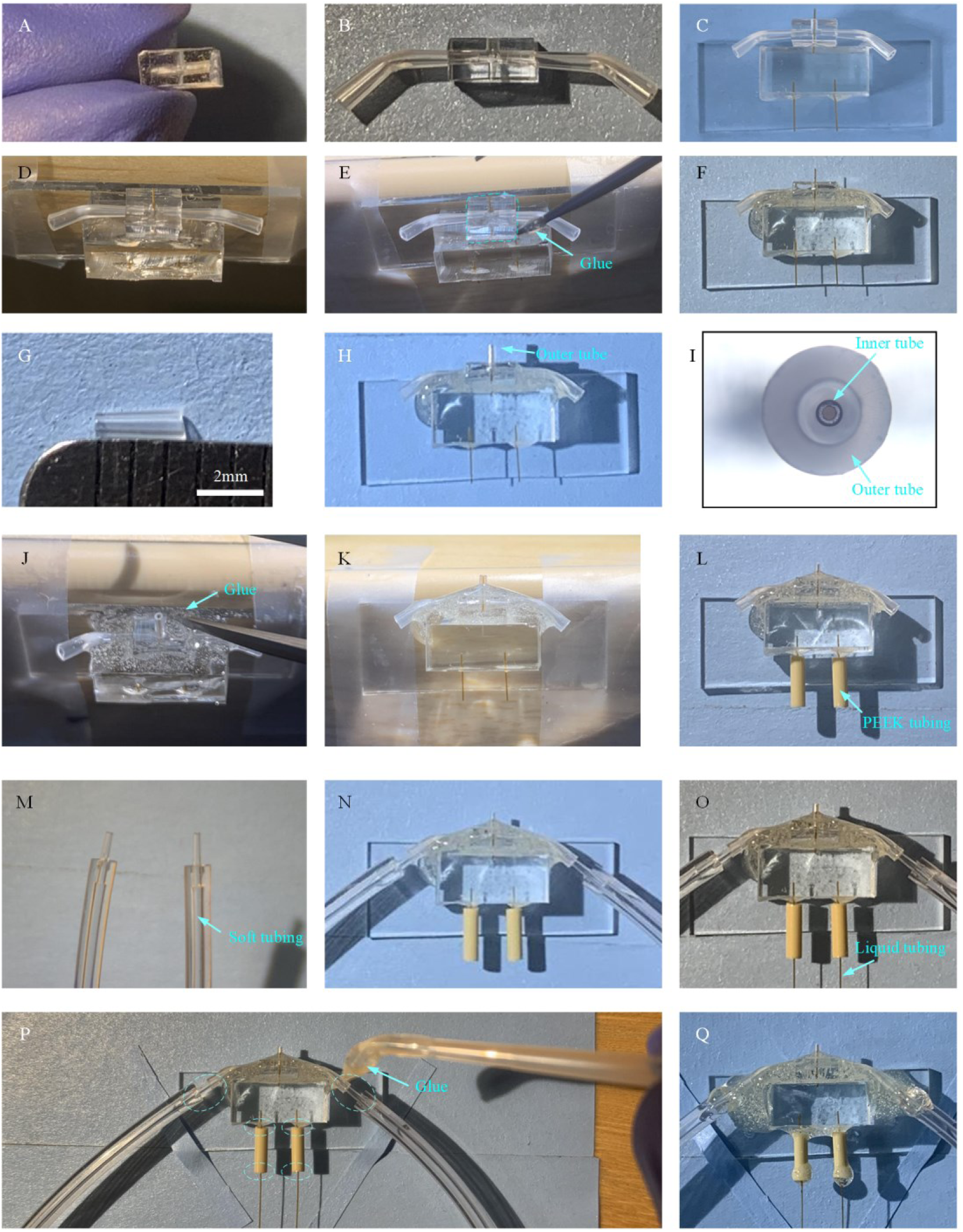
The fabrication of microfluidic chip assembly with time points shorter than 20 ms. A. Prepare smaller nozzle PDMS slab (5 mm x 4 mm x 2 mm) with holes for holding the gas inlet tubings, liquid inlet tubing, and micro-sprayer nozzle. B. Insert gas tubes into the PDMS slab. C. Insert outlet liquid tubing into the back hole of the nozzle PDMS slab. D. Fix the micro-mixer on the side of a table to ensure that the front hole of the nozzle PDMS slab faces up. E. Apply glue to seal the contact area between the micro-mixer and the nozzle PDMS slab. F. The nozzle PDMS glued together with the micromixer. G. Outer tube with a length of 2.5 mm for the microsprayer. H. Insert the outer tube into the front hole of the nozzle PDMS slab to form the microsprayer. I. Outer tube and inner tube (i.e. outlet liquid tubing of the micro-mixer) are aligned to be concentric. J. Fix the micro-mixer on the side of a table to ensure that the outer tube of the micro-sprayer faces up and apply the AB glue to seal the connection between the outer tube and the nozzle PDMS slab. K. The micro-sprayer nozzle with AB glue covered on the corner between the outer tube and the nozzle PDMS slab. L. Put PEEK tubings on the inlets of the micro-mixer. M. Gas tubings. N. Insert the gas tubings into the gas tubes. O. Insert the liquid tubings into the connector PEEK tubings. P. Application of the AB glue to the dash circled areas. Q. Assembled microfluidic chip assembly with time point shorter than 20 ms, ready for use.

**Figure 12.**
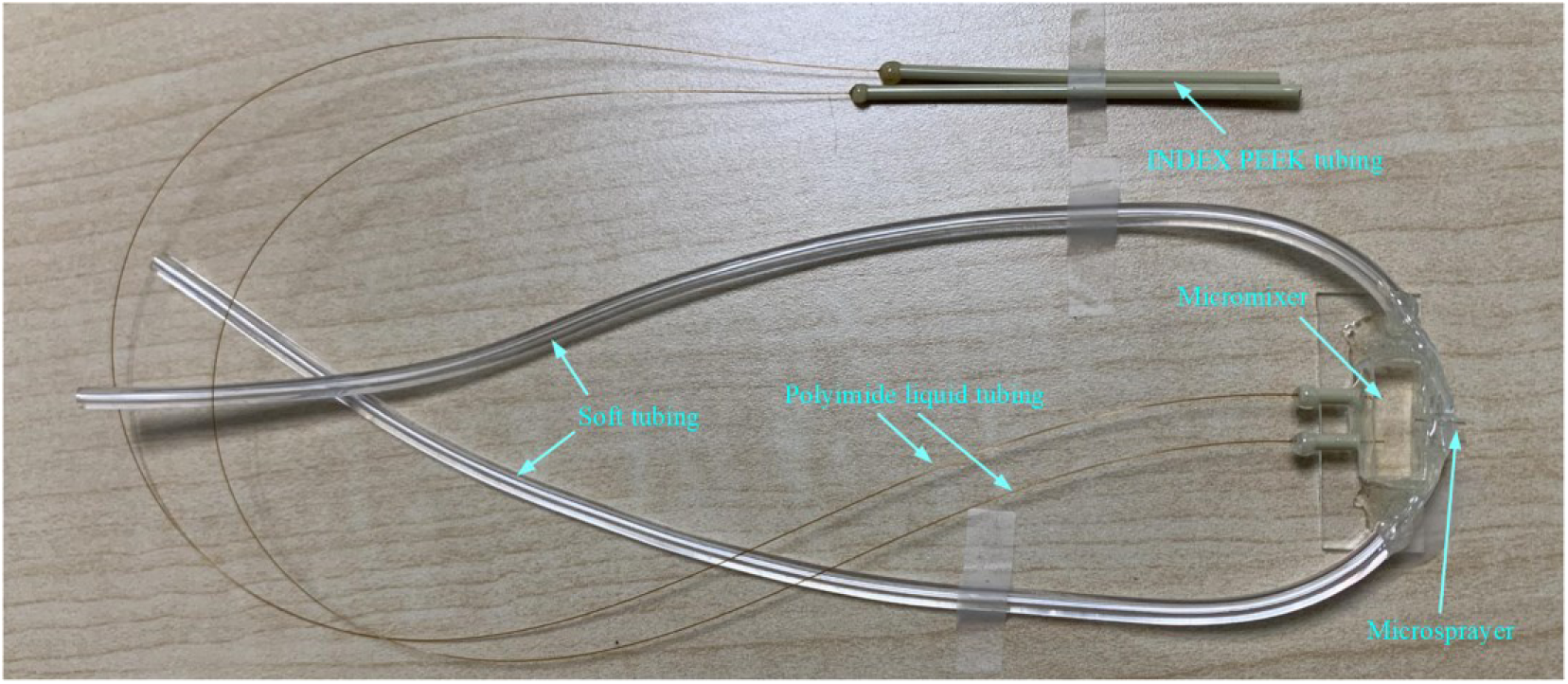
**The entire view of the fabricated microfluidic chip assembly with time points shorter than 20 ms.**

**Figure 13.**
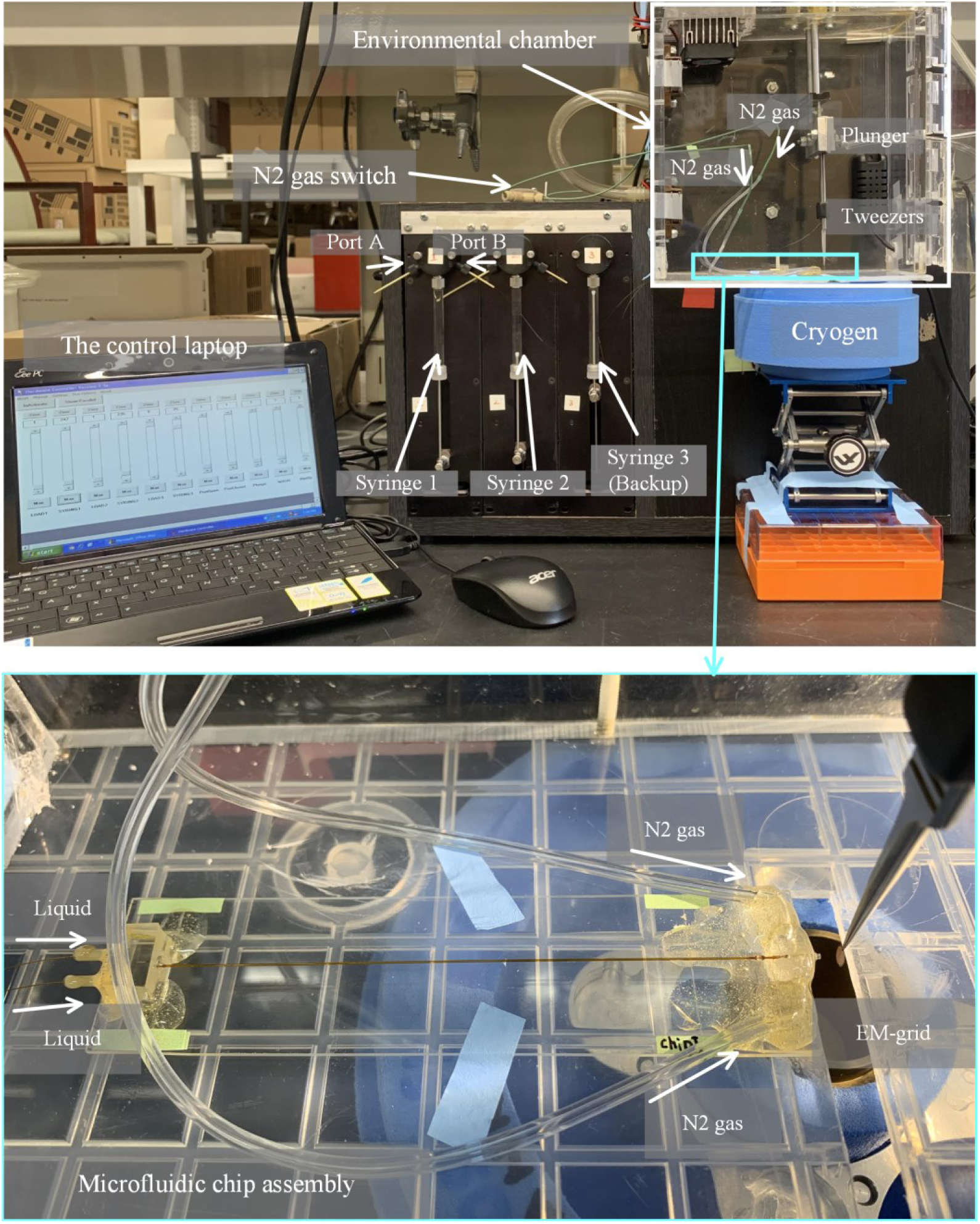
The entire setup with the mounted microfluidic device. (Note that more information can be also seen in Figure S1 and Video S1 in our previous work.[11])

For this protocol, four chip assemblies are fabricated, and time points, and all the key materials and the parameters about the chip assemblies are listed in Table 1. Based on the above-mentioned equation, the reaction times using our chip assemblies could be achieved at 10, 25, 141, and 899 ms, respectively (To be simple, we round them to 10, 25, 140, and 900 ms).

**Table 1.** Materials and parameters for the fabrication of the four chips used in this TR study.

| Microfluidic Chips | Capillary tubing (as microreactor) inner diameter (I.D.) and length ($\mu\text{m}/\text{mm}$) | Tubing (between micromixer and microreactor) I.D. and length ($\mu\text{m}/\text{mm}$) | Tubing (between microreactor and sprayer) I.D. and length ($\mu\text{m}/\text{mm}$) | Sprayer-grid distance (mm) | Sprayer-cryogen distance (mm) | Plunging velocity (m/s) |
| --- | --- | --- | --- | --- | --- | --- |
| Chip 1 (10 ms) | 75/5.0 | N/A | N/A | 3.5 | 10 | 1.9 |
| Chip 2 (25 ms) | 200/2.0 | 75/2.8 | 75/5.0 | 3.5 | 15 | 1.9 |
| Chip 3 (140 ms) | 200/24.0 | 75/3.0 | 75/5.4 | 3.5 | 15 | 1.9 |
| Chip 4 (900 ms) | 300/75.0 | 75/2.6 | 75/5.5 | 3.5 | 15 | 1.9 |

Note: this table was reported as table S1 in our previous study[11].

### E. Preparation of sample for TRCEM

Before the preparation of sample for TRCEM, we would highly recommend doing non-EM kinetic experiments which will save much effort and resources. For this study, as detailed in our previous work[11], we chose different reaction time points according to our peer researchers’ non-EM kinetic results[16], where they followed the kinetics of ribosome dissociation with HflX in stopped flow, by using light scattering as a reporter. In recent years, single-molecule FRET (smFRET) has been used to capture the dynamic process of a reaction system by measuring distances between fluorescence tags within single molecules even at nanometer scale [17], and results from such experiments can guide the choice of time points in the TR experiment, as well.

The entire setup for time-resolved grid preparation, based on the computer-operated liquid-pumping and grid plunging apparatus developed by Howard White (Eastern Virginia Medical School)[18], is depicted in Figure 13. As the key component of the TRCEM apparatus, the microfluidic chip assembly is mounted next to the pneumatic plunger (Figure 13), which are both placed in an environmental chamber[19] that maintains the temperature and humidity. The plunger, which is pneumatically driven and controlled, holds the tweezers on which the EM grid is mounted for fast plunging into liquid ethane after passing the spray cone. In addition, the apparatus contains the pumping system for introducing the solutions into the micromixer and the nitrogen gas into the gas inlets of the microsprayer. Finally, it also includes the computer for controlling both the pumping system and the plunger.

Before each spraying experiment, the tweezers are adjusted in a way that the grid held at their tip passes the nozzle, and then plunges into the cryogen cup. In all our experiments, the condition inside the chamber was maintained at 80%–90% in relative humidity by a humidifier connected to the chamber, and kept at a temperature in the range of 24°C–26°C. Compressed nitrogen gas, humidified by passing through two consecutive water tanks, is fed into the microsprayer at a manually-regulated gas pressure. Once the gas flow is stable, the solutions are injected into the microfluidic chip assembly by syringe pumps under computer control, and the total liquid flow rate is set at 6 µL/s. At this point the sprayer starts spraying. Lastly, the EM grid passes through the spray cone and plunged into liquid ethane. In detail, the procedure of the experiment is as follows:

1. Open the environmental chamber and put the microfluidic chip assembly onto the supporting platform.
2. Mount the tweezers on the plunger and move the plunger up and down to check if the tip of the tweezers will pass through in close vicinity of the opening of the microsprayer nozzle.
3. Align the sprayer nozzle with the tip of the tweezers and move the chip assembly forward and backward (i.e., left and right inside the environmental chamber on Figure 13) to adjust the distance between the sprayer nozzle and the tip of the tweezers to 3.5 mm. After alignment, fix the position of the microfluidic chip using pieces of tape (Fisherbrand™ Labeling Tape).
4. Connect the chip assembly to the syringes 1 and 2 with the tubing (O.D.:150 µm; I.D.: 75 µm) for feeding the sample solution (Figure 13). (Note: the narrow tubing (O.D.:150 µm; I.D.: 75 µm) is used to reduce the dead volume of the sample).
5. Connect the chip assembly with the nitrogen gas tank using the tubing (O.D.: 2.4 mm and I.D.: 0.8 mm) for feeding the gas. There is a gas valve between the gas tank and the microsprayer.
6. Load DI water into the syringes. (As shown in Figure 13, the ports A and B for each syringe valve can be used for withdrawing and dispensing liquid, respectively).
7. Run the syringe pumps for 3 rounds of DI to clean the system. Each time the total volume to be dispensed is set at 20 µL.
8. Turn on the N_2_ gas and at same time run the syringe pumps with DI water. Each time the total volume to be dispensed is set at 20 µL. Increase the gas pressure from 4 to 16 psi and check if the microsprayer keeps working in good condition as shown in Figure 14. (Notes: 1--normally we use 8 or 12 psi as working gas pressure; 2--a laser pointer pen is used to check the spray cone).
9. Discharge the DI water from the syringes, load buffer solution into them, and repeat steps 7 and 8. Check whether the microsprayer works using buffer solution. (Note: check each connection area on the microfluidic chip assembly for possible leakage. It is wise in the planning of an experiment to have one extra chip for backup).
10. Turn on the N_2_ gas tank, which is connected to the plunger. We set the gas pressure for operation of the plunging at 40 psi.
11. Observe and check if the plunger is driven by the N_2_ gas by turning the plunger valve on and off three times using the laptop computer interface.
12. Discharge the buffer solution from the syringes and load two reactant solutions into the empty syringes.
13. Clean the EM-grid with the easiGlow machine to make the grid hydrophilic (15 mA, 35s, and vacuum pressure: 0.26m Torr.)
14. Hold the EM grid by the tweezers.
15. Mount the tweezers on the plunger.
16. Align the EM grid with the sprayer nozzle and adjust the distance in horizontal direction from the nozzle to the EM grid.
17. Turn on the humidifier to increase the humidity inside the environmental chamber and wait for the humidity to stabilize at around 90%.
18. Turn on the N_2_ gas for the plunger and the N_2_ gas for the microsprayer.
19. Move the container with the liquid ethane into the correct position beneath the sprayer nozzle.
20. Turn on the syringe pump and activate the microsprayer to spray. After waiting for the spray plume to stabilize, activate the plunger by the computer, which causes the EM grid to pass through the spray cone and rapidly immerse into the liquid ethane for fast vitrification. (Note that in our experiments we wait 3s to allow stabilization of the spray)
21. Turn off the nitrogen gasses and the humidifier.
22. Unmount the tweezers from the plunger and move the EM grid into a grid box in liquid nitrogen. (Note that for the action of transferring the grid from liquid ethane to liquid nitrogen, the faster the better. As in the blotting method, a lot of practicing is needed for mastering fast transfer.)
23. Repeat steps 13-22 to prepare additional grids using the same microfluidic chip assembly. We normally prepare four grids for use with one chip of a specific reaction time point.
24. Unmount the microfluidic chip assembly and mount a new one with a different reaction time point, then follow the same procedure described above.
25. (Notes: 1–both prior and subsequent to a TR experiment, the microfluidic chip assembly needs to be pumped with DI water for cleaning. 2–Buffer solution is always pre-sprayed to ensure the sprayer chip is in a good working state. 3–Each microfluidic chip assembly is used for one given biological reaction only to avoid cross contamination.)

**Figure 14.**
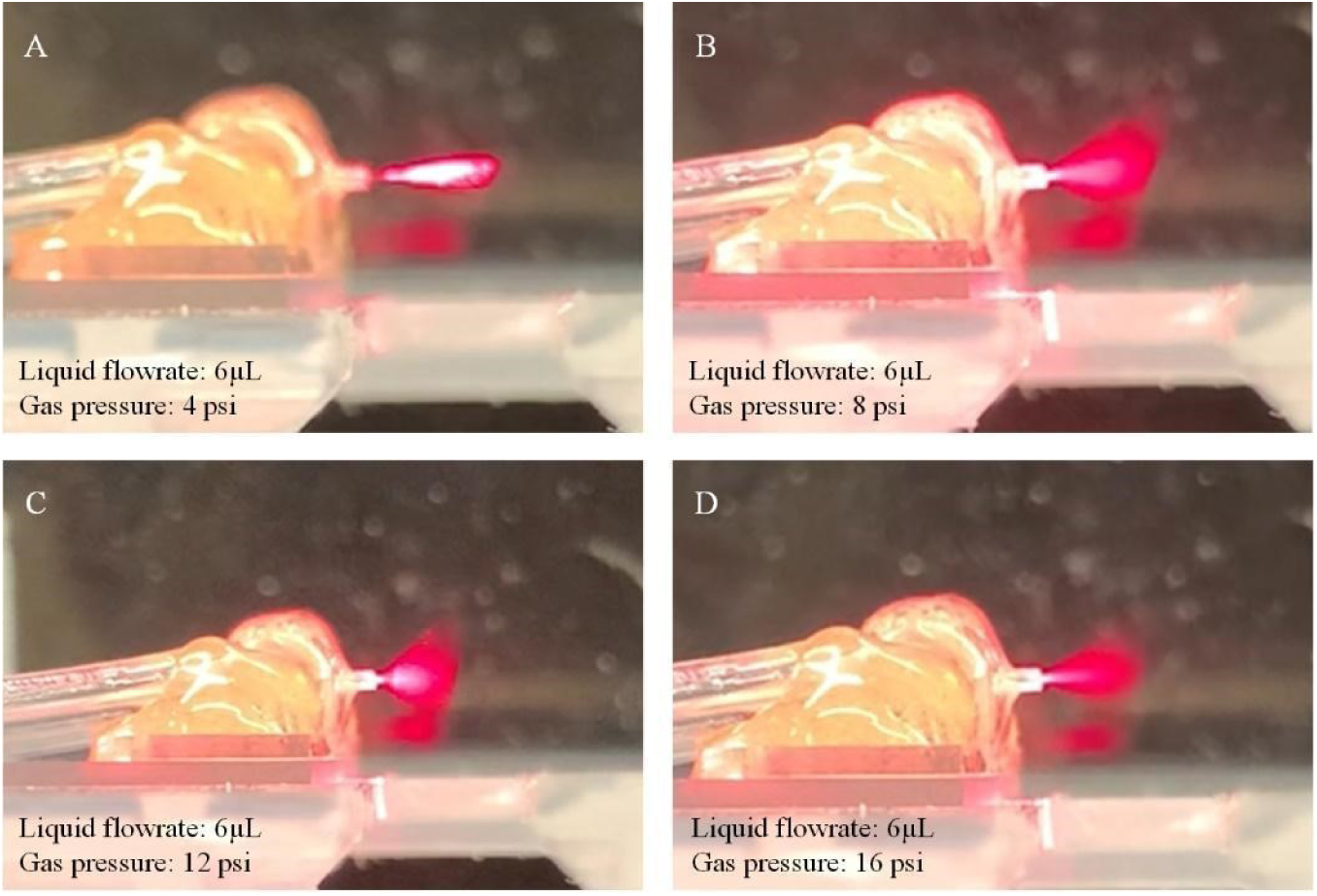
Testing of the entire setup with the microfluidic device. When increasing the gas pressure from 4 to 16 psi (from A to D), good spray performace can be found at 8 and 12 psi. (Notes that a laser pointer pen is used to check the spray cone)

## Data analysis

The data analysis method we employ already appears in sufficient detail in our published paper[11], so a brief summary suffices here. Four respective reactions of 10, 25, 140, and 900 ms were prepared via the TRCEM experimental procedure, and 3452, 3598, 3530, and 3603 good micrographs were collected, respectively, using a commercial high-resolution transmission electron microscope. We always pool together all micrographs from an entire series of TRCEM experiments for data processing (see ref. [3]). After correction of beam-induced motion using MotionCor2 [20] and estimation of the contrast transfer function (CTF) for each micrograph using CTFFIND4 [21] in Relion-4.0 [22], particle picking was performed using Topaz [23]. The entire dataset for the four time points contained around 1 million “good” particles. After further rounds of 2D classification, 802,562 particles were selected, which were used to generate a 3D initial model for 3D auto-refinement in Relion-4.0. CTF refinements were done to correct for magnification anisotropy, fourth-order aberrations, per-particle defocus, and per-particle astigmatism, followed by another 3D auto-refinement. Then 3D classification was performed on the entire pooled dataset without re-alignment, using the angular information from the previous 3D auto-refinement. Seven different classes were found: (1) rotated 70S lacking HflX (r70SnoHflX, 56,894 particles); (2) non-rotated 70S ribosomes lacking HflX (nr70SnoHflX, 96,898 particles); (3) 70S-like intermediate-I bound with HflX (i70SHflX-I, 140,682 particles); (4) 70S-like intermediate-II bound with HflX (i70SHflX-II, 138,296 particles); (5) 70S-like intermediate-III bound with HflX (i70SHflX-III, 113,038 particles); (6) 50S subunit bound with HflX (50SHflX, 62,558 particles); and (7) 30S subunit (58,952 particles) with total of 667,318 particles. For this pooled dataset, a mixture of particle populations with a reaction time range of 10 ms – 900 ms co-exists in each class, and each picked particle was labeled by the time point of the micrograph it originated from, so percentages of particles of different time point contributing to each class can be calculated with respect to the total of 667,318. The detailed description of the analysis can be found in the Cryo-EM data processing Section of the Method Details, or Figure S4D in our work[11].

## Validation of protocol

[This protocol or parts of it has been used and validated in the following research article(s)]:

● Authors Bhattacharjee et al. (2024). Time resolution in cryo-EM using a PDMS-based microfluidic chip assembly and its application to the study of HflX-mediated ribosome recycling. Cell [11] (Figure 1; Figure 2; Figures S1-S2; Figure 3 panel M).

We stored the prepared grid in a liquid nitrogen dewar for future imaging. In this study, after the screening check, we obtained around 2 grids on average for each time point. For data collection, we used 3, 1, 2, 1, and 1 grid(s) for 10 ms, 25 ms, 140 ms, 900 ms, and control, respectively.

All the data were collected using a 300-kV Titan Krios (Thermo Fisher Scientific, Waltham, MA) equipped with a K3 direct detector camera (Gatan, Pleasanton, CA). In contrast to normal blotted grids, the droplet-covered TRCEM grids require manual hole and exposure targeting to pick up collectible droplets and avoid thick-ice areas based on the intensity in the hole images, as shown in the left and middle column panels of Figure 15. Some example high-resolution micrographs from TR grids are shown in the right-column panels of Figure 15. All other details about the data collection can be found in the article referenced above[11]. As a result, we were able to collect 3452, 3598, 3530, and 3603 high-resolution micrographs for 10, 25, 140, and 900 ms, respectively, sufficient for investigating the time course of HflX-mediated ribosome recycling.

**Figure 15.**
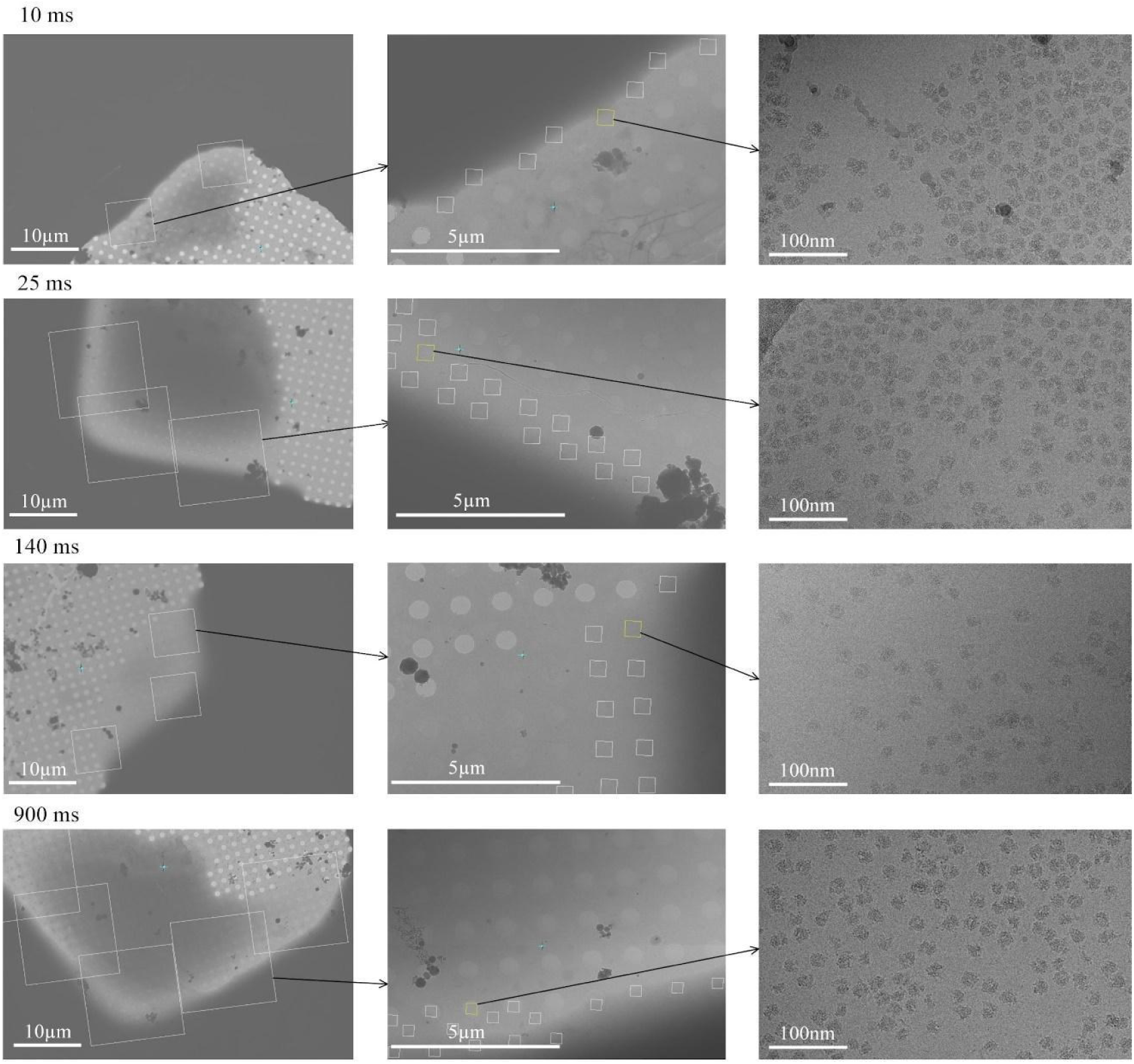
Data collection. The left column panels are micrographs for hole targeting; the middle column panels are micrographs for exposure targeting; and the left column panels are the desired high-resolution micrographs.

As a result of the data processing, we found that the HflX in the presence of GTP interacts with the 70S ribosome and recycles it by splitting it into its 30S and 50S subunits. Three intermediates (Figure 16 A-C) whose occupancies are short-lived yet stretch over two or three time points (Figure 16D) show a stepwise clam-like opening of the 70S ribosome, which allowed us to infer the temporal sequence in which the intersubunit bridges are disrupted. As the density maps were resolved at resolutions around 3 Å, we could build atomic models, allowing the detailed molecular action mechanism of HflX to be inferred. The results demonstrate that this protocol is well adapted to resolve a biological reaction process in the stated time range of 10 to 1000 ms. For more information about the biological story, we refer to the original article [11].

**Figure 16.**
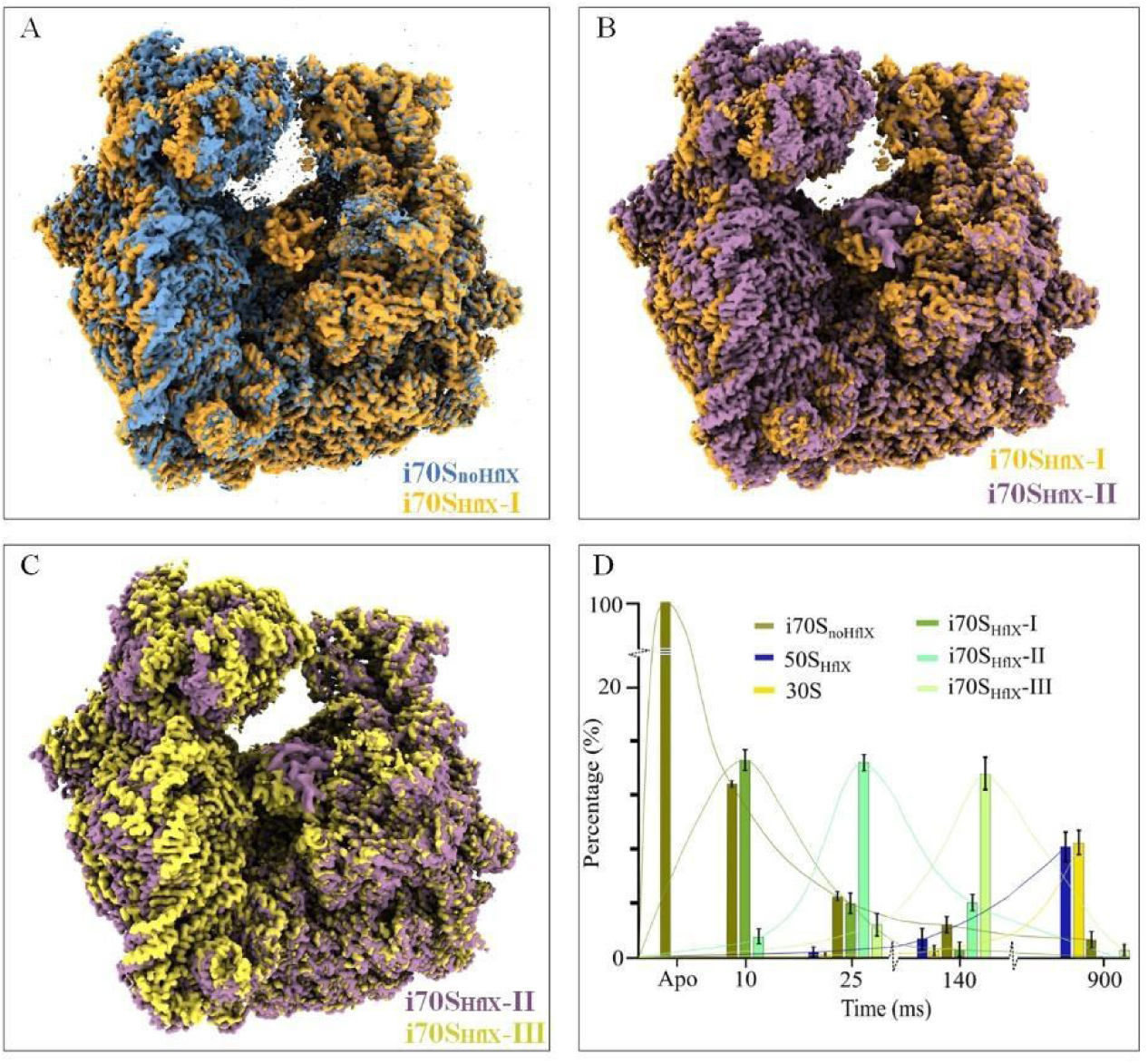
Validation of the protocol. A. Initial opening of the 70S ribosome from i70S_noHflX_ (blue) to i70S_HflX_-(brown), forced by the interaction with HflX. B, C. Motion of the 30S subunit from i70S_HflX_-I (brown) to i70S_HflX_-and to i70S_HflX_-III (yellow), as the 70S ribosome opens stepwise by the action of HflX. The maps of i70S_noHflX_, i70S_HflX_-I, i70S_HflX_-II and i70S_HflX_-III are deposited in the public database as EMD-29681, EMD-29688, EMD-29687 and EMD-29689, respectively. In (A) through (C), all reconstructions are aligned on the 50S subunit. D. Kinetics of the splitting reaction in terms of the number of particles per class (or occupancy of the corresponding state) as a function of time, obtained by 3D classification. Figure 16D was reported as Figure 2M in ref.[11]

## General notes and troubleshooting

### General notes

1. Over the course of three months, we tested the fabricated microfluidic chip assembly and found it can be reused multiple times to prepare TRCEM grids if it is properly cleaned and preserved after each experiment.
2. During the microfabrication, the volume of the PDMS AB mixture used for curing may vary and thus the thickness of the PDMS slab for the sprayer may vary from case to case, and therefore the distance between the sprayer nozzle and ethane cup needs to be measured for each new chip assembly to estimate the plunging time.
3. Copper quantifoil EM grids are preferred as they withstand the force of the spray while gold grids tend to be bent.

### Troubleshooting

Problem 1: Compared with conventional blotting method, the TRCEM method requires a substantially bigger volume of sample (10 µL versus 3 µL) for each grid

Possible cause: (1) “dead volume”: there is a minimum quantity of fluid required to reside in the whole microfluidic system for ensuring a stabilized spray; (2) much of the spray is wasted in the present setup with a single EM grid as target. deposition of the sample from the spray plume onto the grid is inefficient and a lot of the material is wasted.

Solution:While the dead volume problem is difficult to overcome, the waste of active spray can be mitigated by the development of a plunger with multiple pairs of tweezers, or a specially designed tweezer manifold, to hold several grids at once such that they pass successively through the plume.

Problem 2: Compared with the conventional blotting method, the TRCEM method requires a larger volume of sample (10 µL) for each grid.

Possible cause: The tubings to connect the chip and pump syringe need to be filled with the sample and, compared with the conventional blotting method, a larger volume of sample is required to stabilize the spray before the plunging.

Solution: It is essential to develop a plunger with multiple pairs of tweezers or a specially designed tweezer manifold to hold several grids at once.

Problem 3: Data collection on droplet-covered TRCEM grids still relies on manual selection, which is time-consuming.

Likely cause: The droplets have variable sizes and are randomly and often sparsely distributed on the grid surface, and ice thickness can vary from one droplet to another, or even within a droplet (see also Problem 4).

Solution: Deep-learning-based programs may be able to enhance the effectiveness of the data collection on droplet-sprayed EM grids.

Problem 4: Collectible regions of droplets for high-quality images are limited to the narrow regions that touch the grid bar, which limits the amount of data one can obtain from a single EM grid.

Possible cause: the fast plunging and surface tension of the liquid influence the spreading of the droplet on the EM grid.

Solution: Some specialized EM grids, such as self-wicking or nanowire grids [24], ultraflat graphene EM grids [25] are possible options to increase the areas with suitable ice thickness, but these grids need to be strong enough to withstand the force of the gas-assisted spray.

## Supporting information

the Supplementary Mask

## Acknowledgments

This work was supported by a grant from the National Institutes of Health R35GM139453 (to J.F.). All data were collected at the Columbia University Cryo-Electron Microscopy Center (CEC). The microfluidic chips with SiO_2_ coating were fabricated in the nanofabrication clean room facility of Columbia University. We also thank Swastik De for his helpful discussion. The protocol described in this paper is a more detailed account of the experimental setup and experimental results previously published in Cell[11].

## Ethical considerations

No animal and/or human subjects were used in this study.

## References

1. Frank, J. (2017). Time-resolved cryo-electron microscopy: Recent progress. Journal of Structural Biology 200(3): 303–306. 10.1016/j.jsb.2017.06.005.

2. Dandey, V. P., Budell, W. C., Wei, H., Bobe, D., Maruthi, K., Kopylov, M., Eng, E. T., Kahn, P. A., Hinshaw, J. E., Kundu, N., et al. (2020). Time-resolved cryo-EM using Spotiton. Nature Methods 17(9): 897–900. 10.1038/s41592-020-0925-6.

3. Kaledhonkar, S., Fu, Z., Caban, K., Li, W., Chen, B., Sun, M., Gonzalez, R. L. and Frank, J. (2019). Late steps in bacterial translation initiation visualized using time-resolved cryo-EM. Nature 570(7761): 400–404. 10.1038/s41586-019-1249-5.

4. Klebl, D. P., White, H. D., Sobott, F. and Muench, S. P. (2021). On-grid and in-flow mixing for time-resolved cryo-EM. Acta Crystallographica Section D: Structural Biology 77(10): 1233–1240. 10.1107/S2059798321008810.

5. Mäeots, M.-E., Lee, B., Nans, A., Jeong, S.-G., Esfahani, M. M. N., Ding, S., Smith, D. J., Lee, C.-S., Lee, S. S., Peter, M., et al. (2020). Modular microfluidics enables kinetic insight from time-resolved cryo-EM. Nature Communications 11(1): 3465. 10.1038/s41467-020-17230-4.

6. Kontziampasis, D., Klebl, D. P., Iadanza, M. G., Scarff, C. A., Kopf, F., Sobott, F., Monteiro, D. C. F., Trebbin, M., Muench, S. P. and White, H. D. (2019). A cryo-EM grid preparation device for time-resolved structural studies. IUCrJ 6(6): 1024–1031. 10.1107/S2052252519011345.

7. Berriman, J. and Unwin, N. (1994). Analysis of transient structures by cryo-microscopy combined with rapid mixing of spray droplets. Ultramicroscopy 56(4): 241–252. 10.1016/0304-3991(94)90012-4.

8. Lu, Z., Shaikh, T. R., Barnard, D., Meng, X., Mohamed, H., Yassin, A., Mannella, C. A., Agrawal, R. K., Lu, T.-M. and Wagenknecht, T. (2009). Monolithic microfluidic mixing–spraying devices for time-resolved cryo-electron microscopy. Journal of Structural Biology 168(3): 388–395. 10.1016/j.jsb.2009.08.004.

9. Lu, Z., Barnard, D., Shaikh, T. R., Meng, X., Mannella, C. A., Yassin, A. S., Agrawal, R. K., Wagenknecht, T. and Lu, T.-M. (2014). Gas-assisted annular microsprayer for sample preparation for time-resolved cryo-electron microscopy. Journal of Micromechanics and Microengineering 24(11): 115001. 10.1088/0960-1317/24/11/115001.

10. Torino, S., Dhurandhar, M., Stroobants, A., Claessens, R. and Efremov, R. G. (2023). Time-resolved cryo-EM using a combination of droplet microfluidics with on-demand jetting. Nature Methods 20(9): 1400–1408. 10.1038/s41592-023-01967-z.

11. Bhattacharjee, S., Feng, X., Maji, S., Dadhwal, P., Zhang, Z., Brown, Z. P. and Frank, J. (2024). Time resolution in cryo-EM using a PDMS-based microfluidic chip assembly and its application to the study of HflX-mediated ribosome recycling. Cell 187(3): 782–796.e723. 10.1016/j.cell.2023.12.027.

12. Feng, X., Fu, Z., Kaledhonkar, S., Jia, Y., Shah, B., Jin, A., Liu, Z., Sun, M., Chen, B., Grassucci, R. A., et al. (2017). A Fast and Effective Microfluidic Spraying-Plunging Method for High-Resolution Single-Particle Cryo-EM. Structure 25(4): 663–670.e663. 10.1016/j.str.2017.02.005.

13. Schmidt, H. G. (2022). Safe Piranhas: A Review of Methods and Protocols. ACS Chemical Health & Safety 29(1): 54–61. 10.1021/acs.chas.1c00094.

14. Feng, X., Ren, Y. and Jiang, H. (2014). Effect of the crossing-structure sequence on mixing performance within three-dimensional micromixers. Biomicrofluidics 8(3): 034106. 10.1063/1.4881275.

15. Feng, X., Ren, Y. and Jiang, H. (2013). An effective splitting-and-recombination micromixer with self-rotated contact surface for wide Reynolds number range applications. Biomicrofluidics 7(5): 054121. 10.1063/1.4827598.

16. Zhang, Y., Mandava, C. S., Cao, W., Li, X., Zhang, D., Li, N., Zhang, Y., Zhang, X., Qin, Y., Mi, K., et al. (2015). HflX is a ribosome-splitting factor rescuing stalled ribosomes under stress conditions. Nature Structural & Molecular Biology 22(11): 906–913. 10.1038/nsmb.3103.

17. Gentry, R. C., Ide, N. A., Comunale, V. M., Hartwick, E. W., Kinz-Thompson, C. D. and Gonzalez, R. L. (2023). The mechanism of mRNA activation. bioRxiv: 2023.2011.2015.567265. 10.1101/2023.11.15.567265.

18. White, H. D., Walker, M. L. and Trinick, J. (1998). A Computer-Controlled Spraying-Freezing Apparatus for Millisecond Time-Resolution Electron Cryomicroscopy. Journal of Structural Biology 121(3): 306–313. 10.1006/jsbi.1998.3968.

19. Chen, B., Kaledhonkar, S., Sun, M., Shen, B., Lu, Z., Barnard, D., Lu, T.-M., Gonzalez, Ruben L. and Frank, J. (2015). Structural Dynamics of Ribosome Subunit Association Studied by Mixing-Spraying Time-Resolved Cryogenic Electron Microscopy. Structure 23(6): 1097–1105. 10.1016/j.str.2015.04.007.

20. Zheng, S. Q., Palovcak, E., Armache, J.-P., Verba, K. A., Cheng, Y. and Agard, D. A. (2017). MotionCor2: anisotropic correction of beam-induced motion for improved cryo-electron microscopy. Nature Methods 14(4): 331–332. 10.1038/nmeth.4193.

21. Rohou, A. and Grigorieff, N. (2015). CTFFIND4: Fast and accurate defocus estimation from electron micrographs. Journal of Structural Biology 192(2): 216–221. 10.1016/j.jsb.2015.08.008.

22. Kimanius, D., Dong, L., Sharov, G., Nakane, T. and Scheres, S. H. W. (2021). New tools for automated cryo-EM single-particle analysis in RELION-4.0. Biochemical Journal 478(24): 4169–4185. 10.1042/BCJ20210708.

23. Bepler, T., Morin, A., Rapp, M., Brasch, J., Shapiro, L., Noble, A. J. and Berger, B. (2019). Positive-unlabeled convolutional neural networks for particle picking in cryo-electron micrographs. Nature Methods 16(11): 1153–1160. 10.1038/s41592-019-0575-8.

24. Razinkov, I., Dandey, V. P., Wei, H., Zhang, Z., Melnekoff, D., Rice, W. J., Wigge, C., Potter, C. S. and Carragher, B. (2016). A new method for vitrifying samples for cryoEM. Journal of Structural Biology 195(2): 190–198. 10.1016/j.jsb.2016.06.001.

25. Zheng, L., Liu, N., Gao, X., Zhu, W., Liu, K., Wu, C., Yan, R., Zhang, J., Gao, X., Yao, Y., et al. (2023). Uniform thin ice on ultraflat graphene for high-resolution cryo-EM. Nature Methods 20(1): 123–130. 10.1038/s41592-022-01693-y.

