## Supplementary figures and images for "Time-resolved cryo-EM (TRCEM) sample preparation using a PDMS-based microfluidic chip assembly"

### the Supplementary Mask

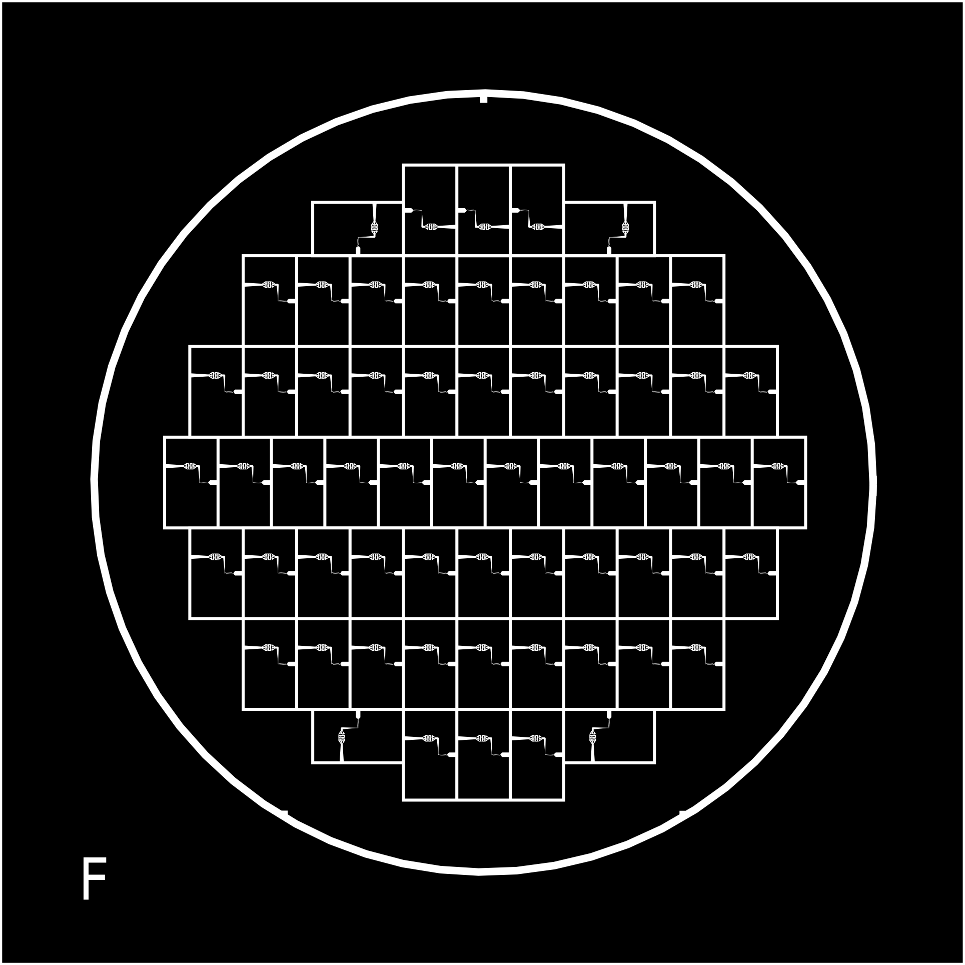
